# Host-cell Interactions of Engineered T cell Micropharmacies

**DOI:** 10.1101/2023.04.05.535717

**Authors:** Christopher M. Bourne, Patrick Wallisch, Megan Dacek, Thomas Gardner, Stephanie Pierre, Kristen Vogt, Broderick C. Corless, Mamadou A. Bah, Jesus Romero Pichardo, Angel Charles, Keifer G. Kurtz, Derek S. Tan, David A. Scheinberg

**Affiliations:** Immunology and Microbial Pathogenesis Program, Weill Cornell Graduate School of Biomedical Sciences, Memorial Sloan Kettering Cancer Center, New York, NY, USA 10065; Molecular Pharmacology Program, Memorial Sloan Kettering Cancer Center, New York, NY, USA 10065; Pharmacology Program, Weill Cornell Graduate School of Biomedical Sciences, Memorial Sloan Kettering Cancer Center, New York, NY, USA, 10021; Tri-Institutional PhD Program in Chemical Biology, Memorial Sloan Kettering Cancer Center, New York, NY, USA, 10065; Louis V. Gerstner Jr. Graduate School of Biomedical Sciences, Memorial Sloan Kettering Cancer Center, New York, NY 10065, USA; Chemical Biology Program, Sloan Kettering Institute, Memorial Sloan Kettering Cancer Center, New York, NY, USA, 10065

## Abstract

Genetically engineered, cytotoxic, adoptive T cells localize to antigen positive cancer cells inside patients, but tumor heterogeneity and multiple immune escape mechanisms have prevented the eradication of most solid tumor types. More effective, multifunctional engineered T cells are in development to overcome the barriers to the treatment of solid tumors, but the interactions of these highly modified cells with the host are poorly understood. We previously engineered prodrug-activating enzymatic functions into chimeric antigen receptor (CAR) T cells, endowing them with an orthogonal killing mechanism to conventional T-cell cytotoxicity. These drug-delivering cells, termed Synthetic Enzyme-Armed KillER (SEAKER) cells, demonstrated efficacy in mouse lymphoma xenograft models. However, the interactions of an immunocompromised xenograft with such complex engineered T cells are distinct from those in an immunocompetent host, precluding an understanding of how these physiologic processes may affect the therapy. Here, we also expand the repertoire of SEAKER cells to target solid-tumor melanomas in syngeneic mouse models using specific targeting with TCR-engineered T cells. We demonstrate that SEAKER cells localize specifically to tumors, and activate bioactive prodrugs, despite host immune responses. We additionally show that TCR-engineered SEAKER cells are efficacious in immunocompetent hosts, demonstrating that the SEAKER platform is applicable to many adoptive cell therapies.

## Introduction

Adoptive T-cell therapies can localize to and kill antigen-positive cells *in vivo* (1–5). These characteristics have led to success against B cell-derived hematopoietic cancers. CAR-T cell products have thus far been approved for B-cell lymphoma, B-cell leukemia, and multiple myeloma. Solid tumors, however, have remained refractory to such therapies and even most B-cell neoplasms ultimately relapse due to tumor heterogeneity and multiple immune escape mechanisms (6). We and many others are further exploring the therapeutic potential of adoptive T cells that can serve as delivery vehicles for drugs or biologic agents, also known as "targeted micropharmacies", in addition to their intrinsic cytotoxic activity (7–14). More complex, and potentially more effective, engineered cells are in development, but the interactions of these highly modified cells with the host are poorly understood. A more complete characterization of targeted micropharmacies in immunocompetent hosts is critical to the safe clinical translation and further development of multifunctional adoptive T cells for the treatment of solid tumors.

Previously, our groups demonstrated that human T cells can be used in an enzyme–prodrug therapy approach to unmask highly toxic drugs selectively at tumors, thus improving antitumor activity (7). The Synthetic Enzyme-Armed KillER (SEAKER) cell platform was first demonstrated with anti-CD19 CAR-T cells secreting the bacterial enzymes β-lactamase (β-Lac) or carboxypeptidase G2 (CPG2). We showed that these enzymes were delivered by CAR-T cells to lymphomas in xenograft mouse models. Furthermore, systemic administration of prodrugs led to unmasking to form the corresponding parent drugs in the tumor microenvironment, leading to delayed tumor growth and improved survival, without systemic toxicity.

While compelling, these previous findings against human lymphoma in xenograft models may incompletely predict the activities of targeted micropharmacies in solid-tumor models, syngeneic systems, or when using TCR-engineered T cells, rather than CAR T cells. Solid tumor microenvironments exploit multiple immune resistance mechanisms, such as antigen escape or downregulation, HLA downregulation, altered peptide presentation, upregulation of inhibitory immune checkpoint ligands, physical extracellular barriers and metabolic dysregulation to thwart T-cell persistence, cytotoxicity, and cytokine secretion (15–18). As a result, solid tumors present difficult challenges to the success of adoptive T cell therapy. Therefore, understanding how T cell drug delivery vehicles localize and deliver therapeutic cargo to solid tumors and interact with the host immune system is of great interest.

T cells undergo homeostatic proliferation in empty T cell niches, which complicates the study of their kinetics and biodistribution in xenograft mouse models (19,20). How SEAKER cells perform in fully immunocompetent hosts has not been characterized. Moreover, the bacterial origin of the SEAKER enzymes used to date necessitates the evaluation of SEAKER enzyme functionality in immunocompetent hosts for further clinical development. In the present study, we investigated: 1) how the engineering of SEAKER cells would affect their pharmacokinetics and biodistribution *in vivo;* 2) if a novel TCR-cell based SEAKER platform could be used to allow us to target *intracellular* antigens and 3) whether the SEAKER platform could be therapeutically effective in immunocompetent hosts. We characterized these critical features and demonstrated that these cellular systems are functional, slowing tumor progression and resulting in significantly enhanced survival. Taken together, these results highlight the feasibility and effectiveness of adoptive T-cell micropharmacies with CAR-T cells *or* TCR-engineered cells in immunocompetent hosts and against human solid tumors.

## Results

### The SEAKER platform is compatible with primary murine T cells

The SEAKER platform has been shown to be efficacious in human B cell lymphoma models (7). However, these xenograft mouse models using NOD-SCID gamma (NSG) mice lack an endogenous adaptive immune system and NK cells, which may affect SEAKER cell kinetics and persistence. Syngeneic models are useful in understanding the behavior of these complex cells for clinical translation. To understand key issues of kinetics, biodistribution, and immunogenicity of the SEAKER platform in immunocompetent hosts, we developed a syngeneic SEAKER cell system. Gene cassettes that included either the secreted β-lactamase (β-Lac) or carboxypeptidase G2 (CPG2) enzymes upstream of GFP via P2A cleavage sites were designed (Figure 1A, Supplemental Figure 1A). The best signal sequences for each enzyme were determined empirically and used in this study (7). We transduced OT-1 T cells, which recognize a chicken ovalbumin peptide (OVA) presented on H-2Kb. Both β-Lac (Figure 1B) and CPG2 (Supplemental Figure 1B) constructs were successfully transduced into primary murine OT-1 T cells, to provide β-Lac OT-1 SEAKER cells and CPG2 OT-1 SEAKER cells for further evaluation herein. A threshold 25% transduction efficiency was set for all experiments unless otherwise noted.

**Figure 1.**
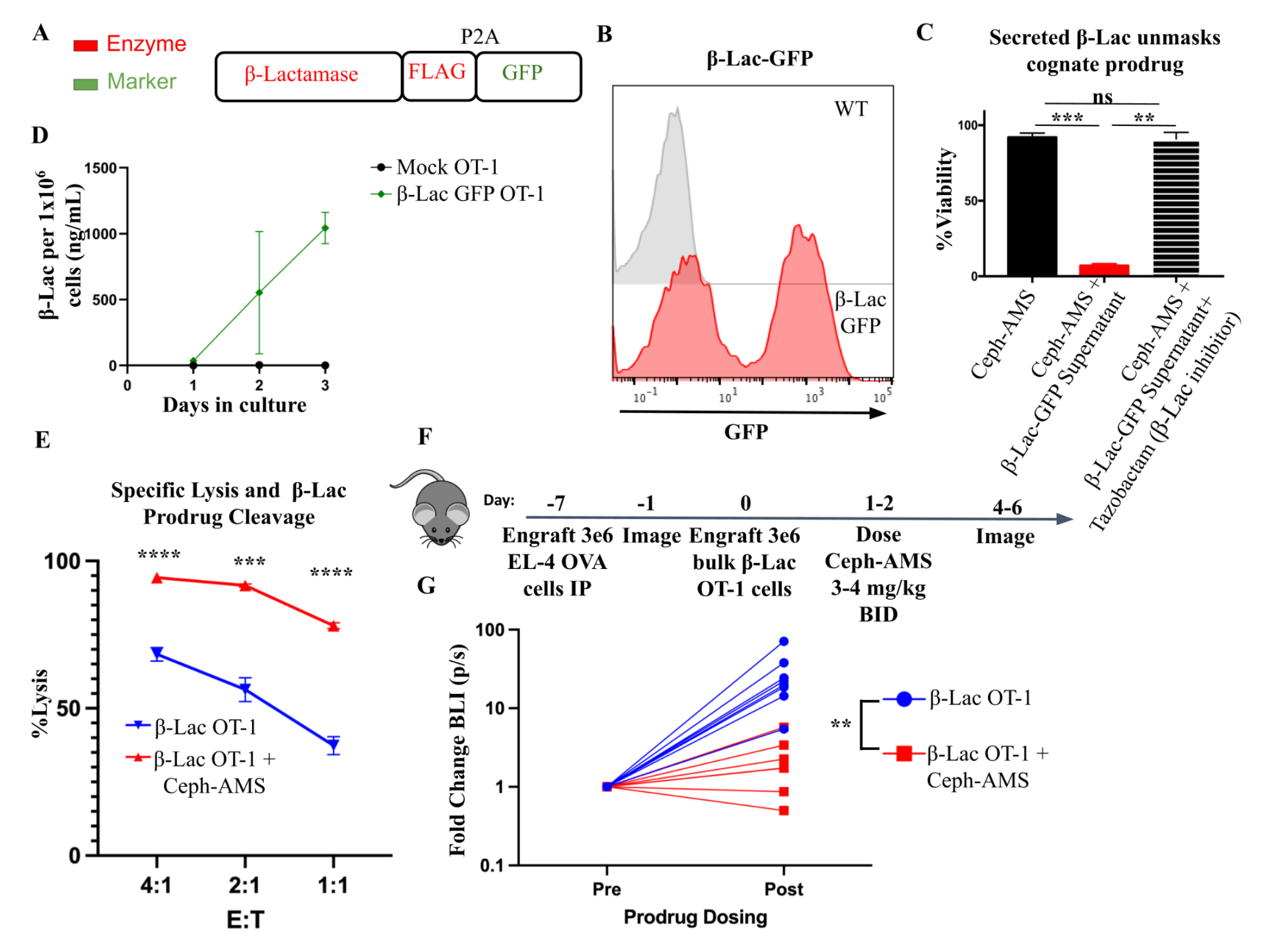
OT-1 syngeneic SEAKER cells secrete functional enzymes and synergize with prodrug to kill cancer cells. (A) Schematic of the SEAKER enzyme secretion cassette for β-lactamase (β-Lac) tagged with flag. (B) Representative flow histogram of murine primary T cell transduction with the β-Lac-GFP plasmid. (C) Supernatant fluid was collected from primary murine T cells transduced with β-Lac-GFP. ffLuc+ EL-4 cells were incubated with 500 nM of the cognate prodrug (Ceph-AMS), with or without the indicated supernatant fluid. β-Lac SEAKER supernatant fluid was also tested with or without tazobactam, a β-Lac inhibitor. Cell viability was assessed by luminescence. (D) Supernatant from β-Lac SEAKER cells was collected and enzyme accumulation was determined by interpolation of values to a standard curve with recombinant β-Lac. (E) Specific lysis of ffLuc+ EL-4-OVA cells with OT-1 T cells expressing β-Lac-GFP. The cognate prodrug Ceph-AMS (500 nM) was added and additional lysis quantified by luminescence readout. (F) Experimental scheme for intraperitoneal proof-of-concept efficacy model. Ffluc+ EL-4 OVA cells were engrafted *IP*. BLI imaging was performed before T cell engraftment IP. Mice were engrafted with 3 x 10^6^ β-Lac OT-1 cells followed by three doses of 3 or 4 mg/kg Ceph-AMS, starting the following day. (G) Mice from both experiments were imaged pre and post prodrug administration and fold change in tumor BLI is graphed. See Supplemental Figure 5 for extended data. Student’s *t* tests were used to determine significance. * = *p*<0.05; ** = *p*<0.01; *** *p*< 0.001; **** = p<0.0001

We have previously demonstrated that the β-Lac enzyme cleaves a Ceph-AMS prodrug to release the potent cytotoxic nucleoside analog AMS (7). Rapidly dividing cells are more susceptible to AMS (7,21). To evaluate the ability of β-Lac OT-1 SEAKER cells to activate Ceph-AMS, spent culture media from these cells was collected and incubated with 180 nM Ceph-AMS and EL-4 tumor cells. While the Ceph-AMS prodrug alone was non-toxic, combination with pre-conditioned media from the murine OT-1 SEAKER cells led to substantial cytotoxicity against the EL-4 target cells, demonstrating that the enzyme was fully functional (Figure 1C).

Addition of the β-Lac inhibitor tazobactam abrogated the cytotoxicity, presumably by blocking conversion of the Ceph-AMS prodrug to the active AMS drug. In the SEAKER system, less than 1 ng/mL of β-Lac can convert non-toxic prodrug Ceph-AMS into highly toxic AMS (Supplemental Figure 2)(7). Using nitrocefin, we were able to quantitate β-Lac secretion at about 500 ng/mL of enzyme per 1 million SEAKER cells, per day (Figure 1D). CPG2 cleaves the AMS-Glu prodrug substrate (7,22). Conditioned media from CPG2 OT-1 SEAKER cells also activated the cytotoxicity of a corresponding AMS-Glu prodrug (100 uM) to induce toxicity on B16F10 cells (Supplemental Figure 1C). Using methotrexate, we were able to quantitate CPG2 secretion at about 200-500 ng/mL per 1 million CPG2-secreting cells (Supplemental Figure 1D,E).

Given the potent cytotoxicity of AMS, we reasoned that the anticancer activity of murine OT-1 SEAKER cells would be improved by the addition of the cognate prodrug. We performed a specific lysis experiment with OT-1 T cells expressing β-Lac co-cultured with the murine T-cell lymphoma cell line, EL-4, ectopically expressing full-length ovalbumin (EL-4 OVA) cells, in the presence or absence of the cognate prodrug (Figure 1E). At each effector-to-target ratio tested, inclusion of the prodrug significantly enhanced cytotoxicity against EL-4 OVA cells. In addition to the OT-1 model, we also generated SEAKER cells based on established CAR-T cell constructs for anti-murine CD19 and anti-MUC16t. Both anti-murine CD19 and anti-MUC16t SEAKER cells secreted enzymes, unmasked prodrug, and displayed enhanced cytotoxicity against antigen-positive and bystander cancer cells (Supplemental Figures 3,4). These data demonstrated the generalizability of the murine SEAKER system to multiple target antigens for CAR (CD19, MUC16t), as well as TCR-modified (OT-1) cells.

### SEAKER cells unmasked prodrugs in a proof-of-concept syngeneic model

As a proof of concept, EL-4 OVA cells were engrafted intraperitoneally (*IP*) in C57BL/6 mice on day –7 to create a fully syngeneic tumor model system to test the SEAKER platform. Baseline tumor volume was measured on day –1, and β-Lac OT-1 SEAKER cells were engrafted *IP* on day 0. Ceph-AMS prodrug administration began on day 1 at either 3 or 4 mg/kg twice a day (*BID*) for three doses (Figure 1F). Mice treated with β-Lac OT-1 SEAKERs and the Ceph-AMS prodrug demonstrated the lowest change in tumor progression across two replicate experiments (Figure 1G, replicates shown in Supplemental Figure 5). These data demonstrated that murine T cells are capable of producing enough enzyme *in vivo* to unmask prodrug and delay tumor growth.

A similar experiment was performed using CPG2 OT-1 SEAKER cells and the corresponding AMS-Glu prodrug (50 mg/kg, BID for 12 doses) (Supplemental Figure 6A). In this experiment, 2 out of 4 mice treated with the combination CPG2 SEAKERs and the AMS-Glu prodrug had a durable response showing no evidence of tumor for weeks (Supplemental Figure 6B-D). The CPG2–AMS-Glu combination showed no overt systemic toxicity in this model (Supplemental Figure 6E) while the β-Lac–Ceph-AMS combination did exhibit significant systemic toxicity and mouse weight loss (Supplemental Figure 5C,H). Because β-Lac has fast enzyme kinetics for conversion of Ceph-AMS into AMS (7), we hypothesized that peritoneal β-Lac caused immediate conversion of the prodrug upon peritoneal injection, leading to systemic leakage of the active AMS drug. On the other hand, CPG2 has slower enzyme kinetics for conversion of its cognate prodrug AMS-Glu (7), and use of this slower system may be beneficial in scenarios where systemic drug leakage is of concern, thus mitigating drug toxicity. Nonetheless, these proof-of-concept experiments demonstrated that both enzyme-prodrug systems had antitumor efficacy in a syngeneic mouse model. However, these experiments utilized *IP* injected SEAKER cells and prodrugs, which simplified the need for specific T cell homing and localization to the tumor.

### Antigen specific SEAKER cells localized to melanoma tumors

Models in which SEAKER cells must traffic and localize to solid tumor masses should better demonstrate the benefits of the SEAKER platform, as compared to peritoneal or *in situ* delivery models. Therefore, to better understand antigen specific SEAKER cell localization in solid tumors, we performed a time course of β-Lac OT-1 SEAKER cell localization to B16F10 melanoma tumors expressing the SIINFEKL ovalbumin peptide (B16-SIIN) in C57BL/6 mice. B16-SIIN tumors were engrafted subcutaneously (*SC*) on day –7.

Cyclophosphamide was injected at 100 mg/kg *IP* to precondition mice for adoptive T cell transfer on day –1. β-Lac OT-1 SEAKERs were engrafted intravenously (*IV*) on day 0. Mice were serially euthanized on days 2, 5, 7, 10 and 14, and tumors and spleens were harvested and analyzed by flow cytometry (Figure 2A). OT-1 T cells have a CD45.1 congenic marker, allowing us to track the total OT-1 T cells as well as the transduced OT-1 SEAKER cells within the same animal. Both untransduced OT-1 T cells and OT-1 SEAKER cells exhibited a peak expansion at day 5 and began contracting by day 7 in the tumor (Figure 2B-E). As anticipated, OT-1 T cells localized at 10-fold lower frequencies in the spleen than tumor, highlighting the specificity of antigen specific T-cell localization. Interestingly, we observed a more pronounced contraction of OT-1 SEAKER cells as compared to untransduced T cells beginning at day 10 (Figure 2F). Nonetheless, the kinetics of the OT-1 SEAKER cells matched that of wild-type OT-1 T cells during the first week of the response, indicating functional fidelity. To further investigate the fitness of β-Lac OT-1 SEAKERs, we compared the tumor control to mock transduced OT-1 T cells at a suboptimal dose of 3×10^6^ cells per mouse. Tumor kinetics were similar between β-Lac OT-1 SEAKERs and mock transduced OT-1 T cells, demonstrating a transient control of tumor as compared to PBS treated mice (Figure 2G,H).

**Figure 2.**
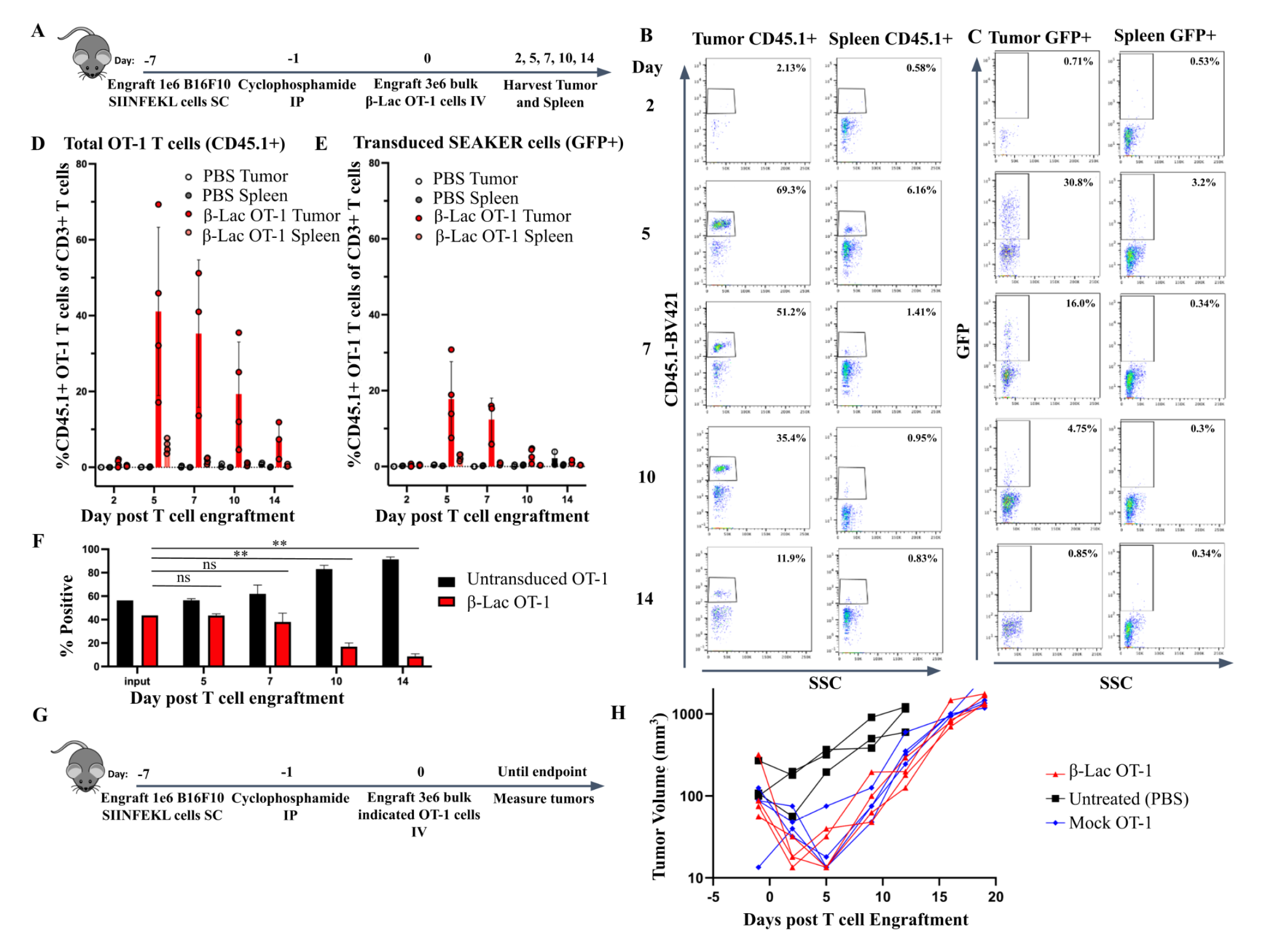
β-Lac OT-1 SEAKER cells localize specifically to B16-SIIN tumors at day 5 post T cell injection. (A) Experimental scheme for time course flow cytometry experiment in which C57BL/6 mice were engrafted with B16-SIIN cells *SC* for 7 days. One day prior to T cell injection, mice were preconditioned with 100 mg/kg cyclophosphamide. On day 0, 3 x 10^6^ OT-1 T cells transduced with β-Lac GFP were engrafted retro-orbitally. Tumors and spleens were harvested at indicated time points and analyzed by flow cytometry. (B) Representative flow plots for CD45.1+ OT-1 T cells in tumor and spleen samples. (C) Representative flow plots for transduced GFP+ SEAKER cells in tumor and spleen samples. (D) Graph of experiment in panel A for CD45.1+ OT-1 T cells. n=2 for PBS, n=3–4 for β-Lac OT-1 per time point. (E) Graph of experiment in panel A for transduced GFP+ SEAKER T cells. (F) The ratio of transduced SEAKER cells to untransduced OT-1 T cells are compared to the input transduction efficiency. (G-H) Mice were treated as in A and GFP-Luciferase (mock) or β-Lac-GFP transduced OT-1 T cells were engrafted *IV*. Tumors were measured using calipers. Student’s *t* tests were used to determine significance. * = *p*<0.05; ** = *p*<0.01; *** *p*< 0.001; **** = *p*<0.0001.

T cells naturally localize to antigen-positive tissues, but also localize to secondary lymphoid organs (23), which might cause off-target toxicity in the SEAKER platform, due to prodrug unmasking. To understand whether SEAKER cells localize to off-target tissues, we treated B16-SIIN-bearing mice and harvested tumors, major secondary lymphoid organs (draining lymph nodes, bone marrow and spleen), blood and the lungs at day 6 post β-Lac OT-1 SEAKER cell engraftment (Figure 3A). As anticipated, the β-Lac OT-1 SEAKER cells localized at high percentages in the tumor, but were present only at very low frequencies in off-target organs (Figure 3B-E). Taken together, these findings demonstrated that SEAKER cells localize specifically to tumors and were rarely found in off-target secondary lymphoid organs during the peak expansion, which indicates a reduced risk of off-target toxicity.

**Figure 3.**
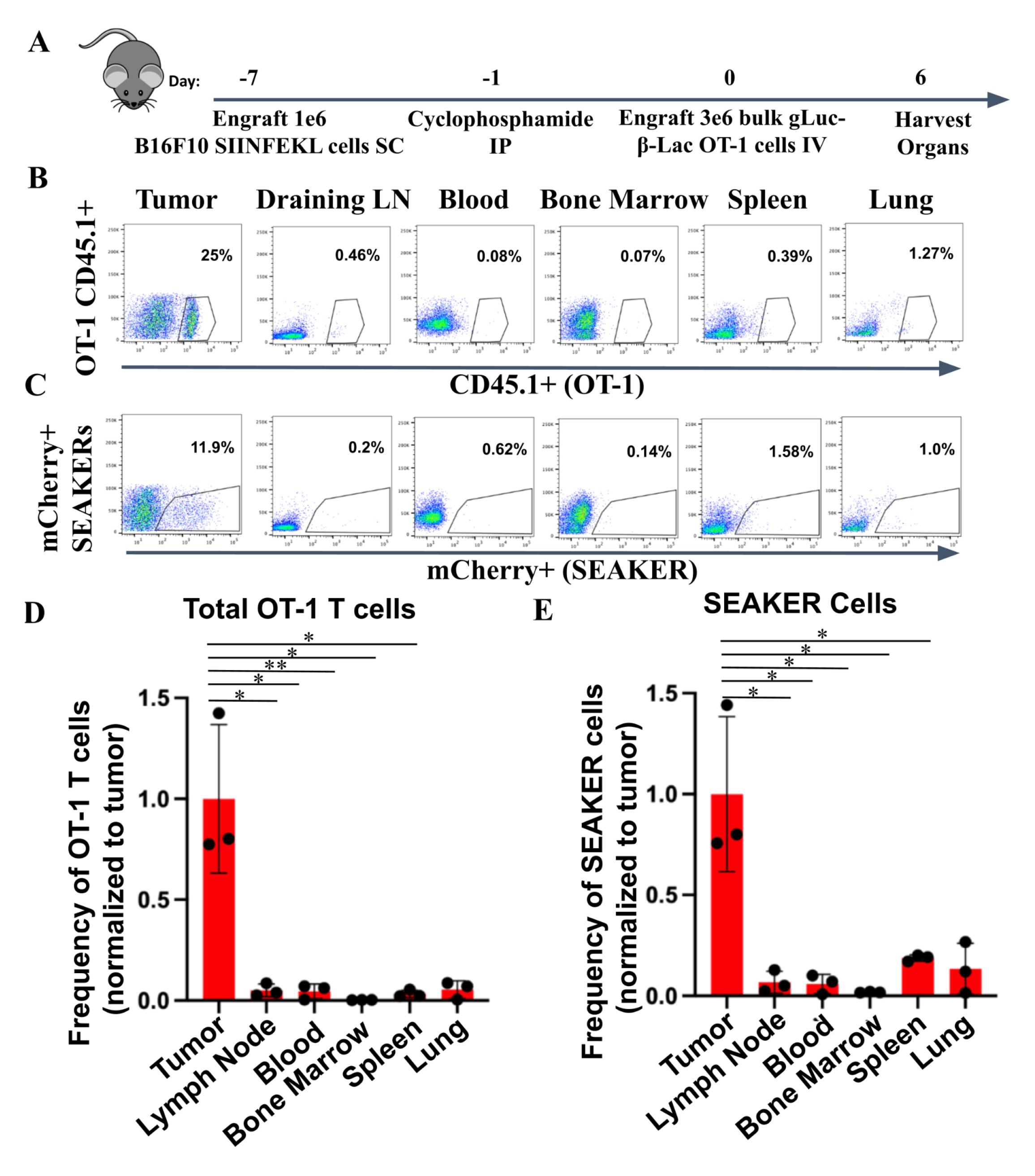
β-Lac OT-1 SEAKER cells localize specifically to tumors and not secondary lymphoid organs. (A) Experimental scheme for standard B16-SIIN tumor model in which indicated organs are harvested on day 6 post T cell engraftment and analyzed for SEAKER cell counts via flow cytometry. (B) Representative flow plots of CD45.1+ OT-1 T cells. (C) Representative flow plots of mCherry+ SEAKER T cells. (D) Graph of CD45.1+ OT-1 T cells in indicated organs, n=3. (E) Graph of mCherry+ SEAKER T cells in indicated organs. Student’s *t* tests were used to determine significance. * = *p*<0.05; ** = *p*<0.01; *** *p*< 0.001; **** = *p*<0.0001

### SEAKER cells deliver functional synthetic enzymes to tumors

To study the pharmacokinetics of SEAKER cells *in vivo*, we used *Gaussia* luciferase (gLuc), which enables the bioluminescent imaging (BLI) of antigen specific T cells in both tumor and healthy tissues (Supplemental Figures 7-9)(24). To track the exact localization of enzyme secreting cells, we designed a β-Lac OT-1 SEAKER construct that includes a membrane-anchored gLuc, and mCherry. These cells were used in the B16-SIIN model described above and mice were serially imaged by BLI (Figure 4A-B). In accordance with the flow cytometric analysis of SEAKER cells (Figure 2), we observed a peak in expansion at day 5 post T cell engraftment followed by a contraction of detectable cells (Figure 4C-D). Administration of rhIL-2 at 4.5 x 10^5^ IU/mouse/day for 6 days to stimulate T cell growth further did not impact SEAKER cell expansion (Figure 4D). Similar expansion kinetics were seen in the analogous CPG2 OT-1 SEAKER models (Supplemental Figure 10). Taken together, these results demonstrated that SEAKER cell kinetics can be tracked longitudinally using BLI. Importantly, two separate methods of kinetic tracking (flow cytometry and BLI) both demonstrated the same kinetics, despite introduction of the foreign gLuc protein.

**Figure 4.**
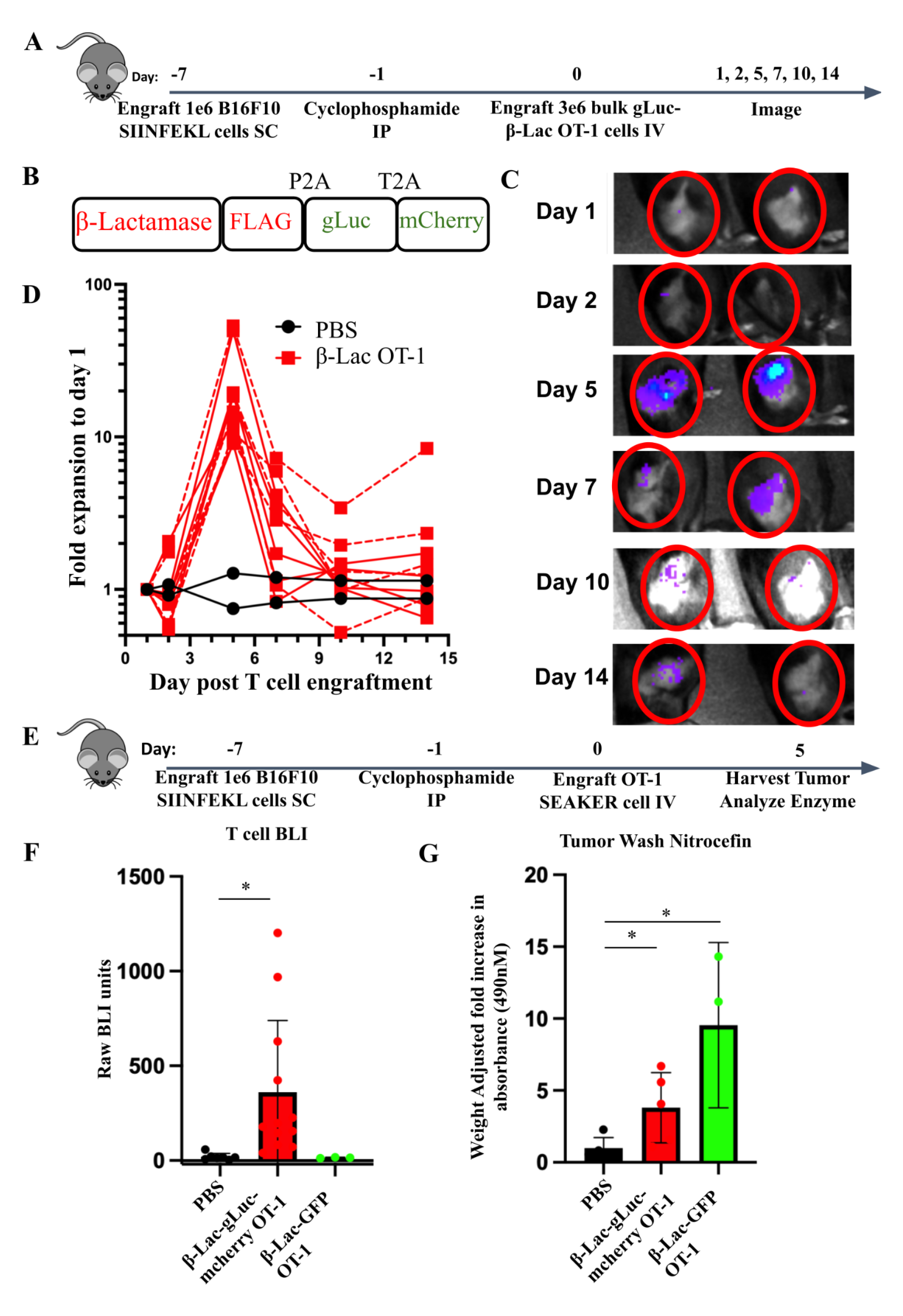
β-Lac OT-1 SEAKER cells deliver enzymes to B16-SIIN tumors at day 5 post T cell injection. (A) Experimental scheme for time course bioluminescent imaging (BLI) of SEAKER cell pharmacokinetics in the B16-SIIN model. (B) Cassette for tracking SEAKER cell localization using membrane-anchored *Gaussia* luciferase (gLuc). (C) Representative images of experiment described in panel A. (D) T cell imaging graph of the experiment described in panel A. Dotted lines represent animals treated with rhIL-2 at 4.5 x 10^5^ IU/mouse/day. n=2 PBS, n=10 β-Lac OT-1. (E) Schematic for enzyme activity in B16-SIIN bearing mice. Tumors were harvested at day 5 post T cell injection and homogenized. (F) Tumor homogenates were mixed with coelenterazine and luminescence was read on a plate reader. PBS n=5, β-Lac-gLuc OT-1 SEAKER, n=12, β-Lac-GFP OT-1 SEAKER, n=3. (G) Cell-free tumor washes were mixed with nitrocefin and enzyme substrate cleavage was measured by absorbance at 490 nm. PBS n=5, β-Lac-gLuc OT-1 SEAKER, n=5, β-Lac-GFP OT-1 SEAKER, n=3. Student’s *t* tests were used to determine significance. * = *p*<0.05; ** = *p*<0.01; *** *p*< 0.001; **** = *p*<0.0001.

Although localization of SEAKER cells to antigen-positive tumors was robust, it was not yet evident whether these cells were delivering functional enzymes to tumors in the immunocompetent hosts. Thus, we harvested tumors from mice treated with β-Lac OT-1 SEAKER cells and extracted the soluble protein fraction (Figure 4E). When treated with the gLuc substrate coelenterazine, only tumor homogenates from the β-Lac-gLuc-mCherry OT-1 SEAKER-treated mice had any detected BLI signal *ex vivo* (Figure 4F). We then mixed tumor homogenate extracts with nitrocefin to assess β-Lac enzyme function directly. Both the tri-cistronic β-Lac-gLuc-mCherry OT-1 SEAKER and bi-cistronic β-Lac-GFP OT-1 SEAKER treated mice had significantly more enzyme activity than control mice in the tumor homogenates (Figure 4G), and the bi-cistronic vector resulted in higher levels of expression.

These results showed that the level of secretion of β-Lac enzyme *in vivo* can be fine-tuned through the number of 2a elements. Each additional 2a site on the vector decreases the production efficiency of each genetic element. Importantly, levels of enzyme activity were sufficient to convert substrate *ex vivo*, indicating the feasibility of the SEAKER–prodrug approach in immunocompetent hosts. We also showed that targeted delivery of β-Lac enzyme was achievable in a peritoneal ovarian tumor model using the anti-MUC16 CAR model (Supplemental Figure 11). This demonstrated that the SEAKER platform can deliver enzymes using both CAR and TCR-based antigen targeting mechanisms to a variety of anatomical locations in immunocompetent hosts.

### SEAKER cells synergize with prodrug to delay melanoma progression

Having established that SEAKER cells are functional *in vivo* and localize specifically to B16-SIIN tumors, we designed an efficacy model to test whether β-Lac OT-1 SEAKER cells can be potentiated with the non-toxic Ceph-AMS prodrug to delay tumor growth. Consistent with a previous report (25), direct treatment with the AMS parent drug induced pronounced systemic toxicity in mice, manifested by rapid weight loss, and had only a modest anti-tumor effect against B16 at the maximum tolerated dose of 0.1 mg/kg (Supplemental Figure 12). To determine whether the SEAKER platform could improve anti-tumor efficacy and decrease toxicity, B16-SIIN tumor-bearing mice in our syngeneic model were engrafted with β-Lac OT-1 SEAKER cells on day 0, then treated with Ceph-AMS on days 4-8 (4 mg/kg, *BID* for 10 doses) (Figure 5A). Mice treated with β-Lac OT-1 SEAKER cells and prodrug demonstrated delayed tumor growth as measured by digital calipers (Figure 5B). Furthermore, survival of mice given SEAKER cells and prodrug was significantly enhanced compared to mice treated with SEAKER cells or prodrug alone (Figure 5C). Importantly, we observed no overt toxicity as evidenced by no decrease in weight of mice treated with SEAKER plus prodrug (Figure 5D). These results showed that the SEAKER platform is efficacious against melanoma solid tumors without the characteristic toxicity associated with direct administration of the AMS parent drug, due to local generation at the tumor.

**Figure 5.**
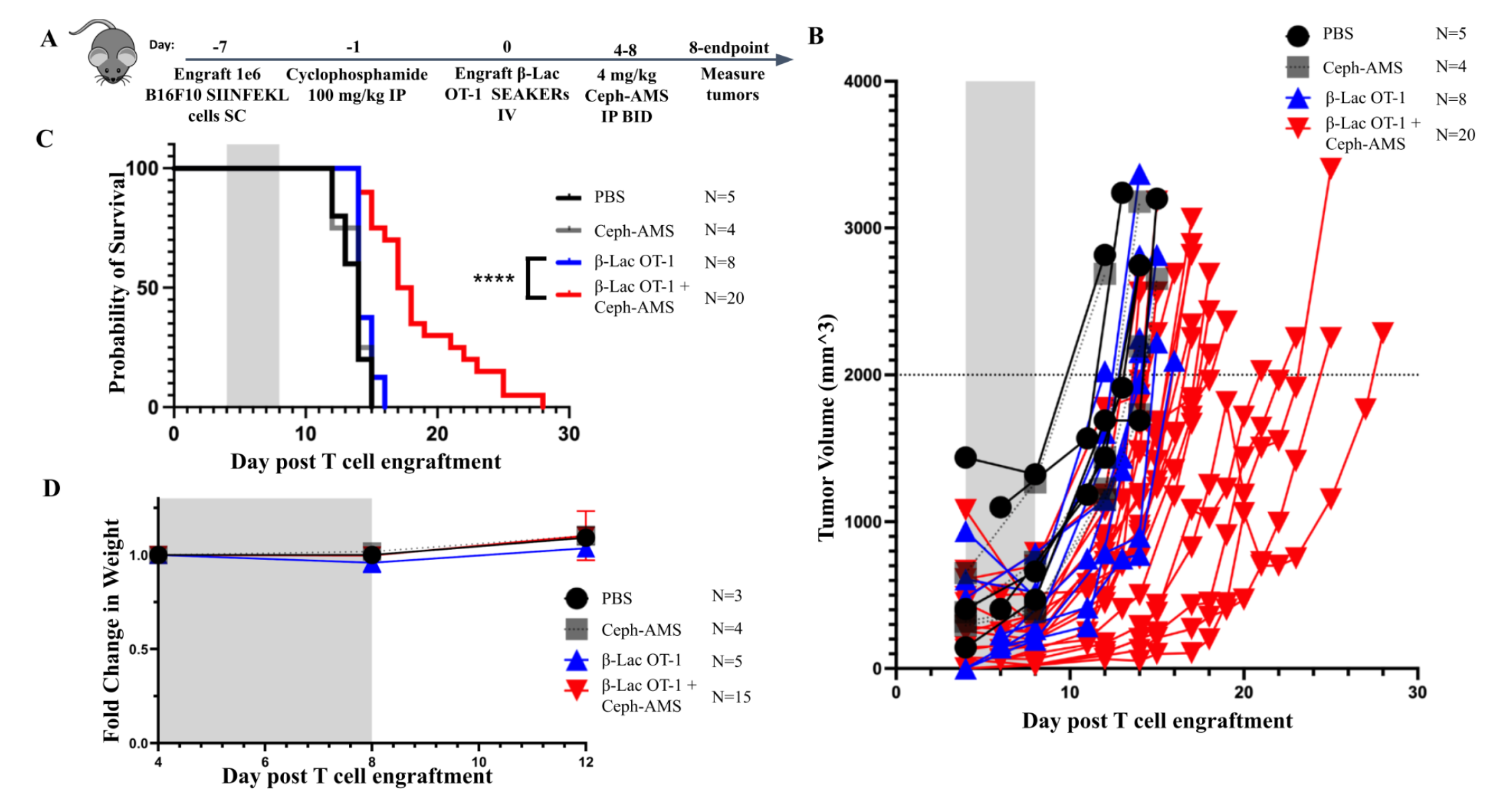
β-Lac OT-1 SEAKER cells unmask Ceph-AMS prodrug to delay tumor growth *in vivo*. (A) Experimental scheme for the B16-SIIN tumor efficacy model. C57BL/6 mice were engrafted with 1 x 10^6^ B16-SIIN tumor cells *SC* and treated with 3 x 10^6^ β-Lac OT-1 SEAKER cells retro-orbitally. On days 4–8, 4 mg/kg of Ceph-AMS was injected *IP, BID*. Tumor volumes were measured by calipers until endpoint. (B) Tumor volumes for mice treated with Ceph-AMS prodrug alone, β-Lac OT-1 SEAKER cells alone, or SEAKER + Ceph-AMS, from 2 repeated experiments. (C) Survival of mice from two repeated efficacy experiments. Significance assessed by log-rank Mantel–Cox test. * = *p*<0.05; ** = *p*<0.01; *** *p*< 0.001; **** = *p*<0.0001. (D) Percent change in weight of mice on day 8 post T cell injection, after prodrug or vehicle administration.

### SEAKER cells survive prodrug treatment in the tumor

One potential limitation of the SEAKER platform is that the SEAKER cells may themselves be subject to the cytotoxic effects of the prodrug they unmask. To assess this issue, we used our trackable SEAKER cells in the *SC* efficacy model established above (Figure 5) and imaged for the presence of the T cells every day during the prodrug dosing regimen (Figure 6A). Surprisingly, groups of mice given the SEAKER cells alone or SEAKERs plus prodrug demonstrated identical T-cell pharmacokinetics during the dosing schedule (Figure 6B-C). We speculate that the SEAKER cells may survive because the prodrug is administered at the end of the T cell expansion period, when the T cells are presumably slowing proliferation. This may explain the unexpected resistance of SEAKER cells to the active cytotoxic drug unmasked *in vivo*.

**Figure 6.**
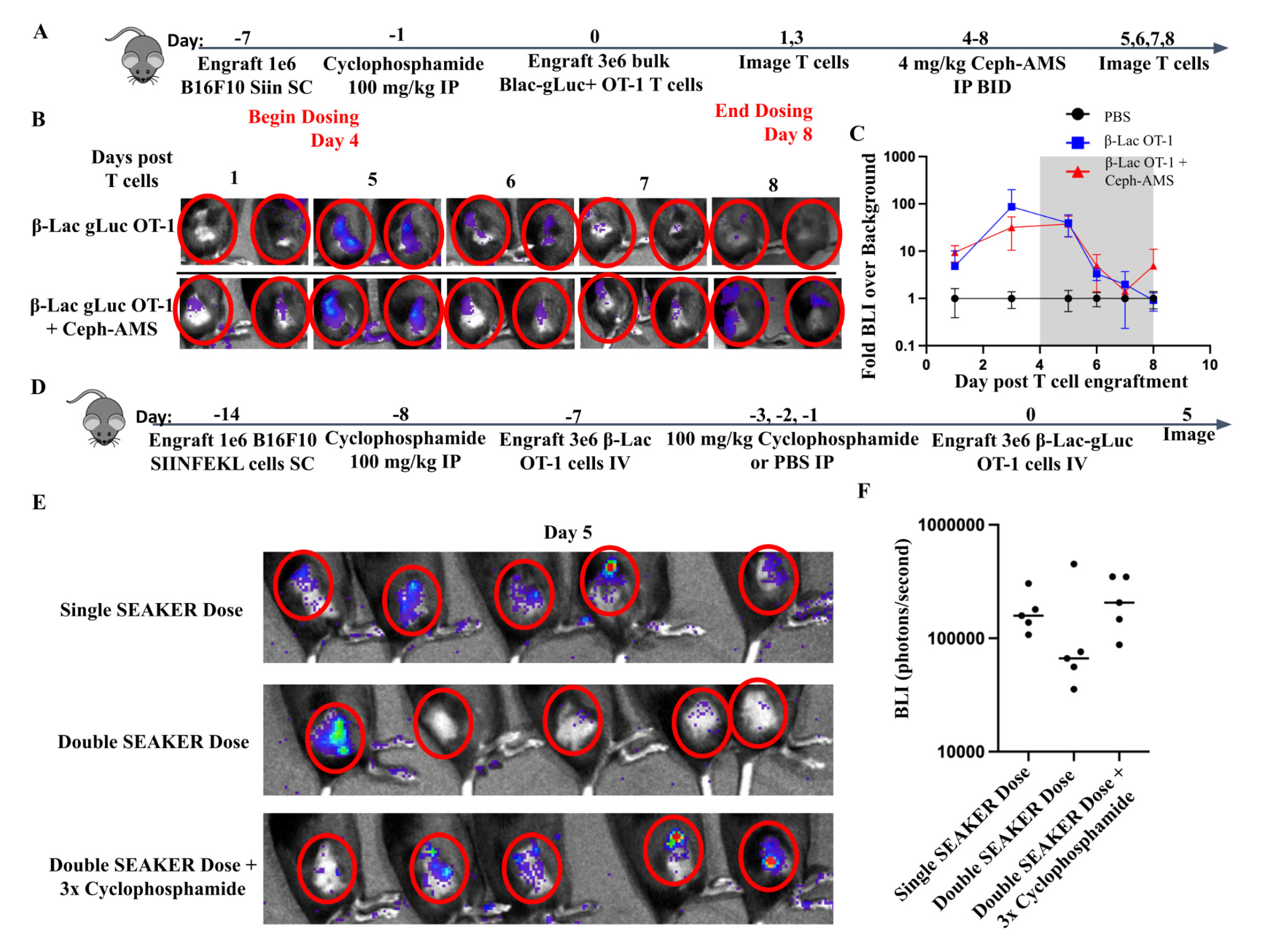
SEAKER cells survive prodrug unmasking and can overcome cellular immune responses. (A) Experimental scheme for T cell trafficking experiment using the B16-SIIN efficacy model. gLuc+ T cells were imaged before and during prodrug administration in the efficacy model. (B) Representative images of T cell tracking in the tumors. Red circles highlight the tumor location. (C) Quantification of time course imaging experiment as described in panel A, PBS N=2, β-Lac OT-1 N=3, and β-Lac OT-1 + Ceph-AMS N=5. (D) Experimental scheme for B16-SIIN model testing repeat engraftment of SEAKER cells. B16-SIIN cells were engrafted *SC* at day –14, pretreated with 100 mg/kg cyclophosphamide *IP* at day –8 and treated with 3 x 10^6^ β-Lac OT-1 SEAKER cells retro-orbitally on day –7. One group of mice received 3 doses of cyclophosphamide on days –3, –2, and –1, while the other two groups received one dose on day –1. Trackable gLuc β-Lac OT-1 SEAKERs were engrafted on day 0 and mice were imaged on day 5. (E) Day 5 images of T cell engraftment in the tumor. (F) Quantification of raw BLI of images in panel B.

### Assessing the immunogenicity of the β-Lac SEAKER cells in immunocompetent hosts

The bacterial origin of SEAKER enzymes raises the possibility that they will be immunogenic in immunocompetent hosts, potentially preventing the cellular or enzymatic functions. Mice treated with β-Lac OT-1 SEAKERs mounted a humoral response to the enzyme that peaked at day 5 post T cell injection, which followed a similar kinetic time course to that of the SEAKER cells themselves (Supplemental Figure 13A-E). Importantly, when those same serum samples were incubated with recombinant β-Lac and nitrocefin substrate, equivalent levels of substrate cleavage resulted from enzymes treated with naive mouse serum or β-Lac OT-1 SEAKER-treated mouse serum (Supplemental Figure 13F-G). This indicates that, although the antibodies that bind β-Lac are induced, they do not block the function of the enzyme.

Multiple rounds of adoptive T cell transfer might increase anti-tumor efficacy and T cell localization in the tumor. However, immune responses against adoptive T cells have been documented after single injections, preventing additional rounds of cell administration (8). We assessed whether SEAKER cells could be readministered in previously-treated mice to explore the impacts of retreatment upon the pharmacokinetics of the cells. B16-SIIN tumor-bearing mice received a preconditioning dose of cyclophosphamide at 100 mg/kg *IP* at day –8 then a dose of either PBS or β-Lac-GFP OT-1 SEAKER cells at day –7. Mice were then stratified into two further groups: one cyclophosphamide dose or three consecutive cyclophosphamide doses at 100 mg/kg *IP*. gLuc+ SEAKER cells were then engrafted into all groups on day 0 and imaged day 5 post engraftment (Figure 6D). Mice that had not been pretreated engrafted the SEAKERs by day 5. In contrast, four of five mice treated with two successive SEAKER doses and one cyclophosphamide dose rejected the second SEAKER cell engraftment (Figure 6E-F). However, mice who received three doses of cyclophosphamide prior to the second adoptive transfer showed enhanced engraftment of the second SEAKER administration (Figure 6E-F). Furthermore, primary engraftment of wild-type OT-1 T cells or CPG2 OT-1 SEAKER cells had no impact on subsequent engraftment of β-Lac OT-1 SEAKER cells (Supplemental Figure 14). These results demonstrated that the host immune responses to SEAKER cells can be overcome through simple pharmacological manipulation. Taken together, these studies show that the SEAKER-produced enzymes are still functional despite being bound by antibodies, and that multiple rounds of SEAKER cell treatment can be achieved with stringent preconditioning, similar to what is used in humans clinically.

## Discussion

We have examined the kinetics, biodistribution, and efficacy of complex, cargo-delivering, adoptive T cells in immunocompetent hosts bearing solid tumors, revealing both opportunities and potential challenges. The results presented demonstrate the feasibility and efficacy of adoptive T cell micropharmacies in syngeneic systems. Traditionally optimized for their cytotoxic capacity, T cells can now be used for consistent, large-scale, highly-localized cargo delivery. This will allow for effective therapeutic killing, even for poorly expressed, downregulated, or heterogenous antigens within a tumor with limited access. The current study presents four conceptual advances for adoptive T cell micropharmacies: 1) cargo delivery and efficacy in an immunocompetent system, 2) cargo delivery and efficacy in solid-tumor models, 3) expansion of the target repertoire of SEAKER cells to intracellular antigens using TCRs, and 4) circumvention of the SEAKER and enzyme immunogenicity issues. Syngeneic tumor models enable the study of adoptive T cells in immunologically competent hosts. In contrast, xenograft models in NSG mice lack common gamma chain-dependent immune cells, such as T cells, B cells, and NK cells (26,27). Furthermore, innate immune function in these models is altered. As a consequence, secondary lymphoid organ formation is also altered. Competition for pro-survival cytokines is largely non-existent, which allows adoptively transferred human cells to maintain homeostatic proliferation in the absence of antigen, confounding pharmacokinetic and biodistribution studies (19,20). Critically, anti-mouse reactive human cells selectively expand in xenograft models, and induce graft-versus-host disease at late timepoints (28). Investigation of true T cell pharmacokinetics is not feasible in these models. Syngeneic mouse models circumvent all of the caveats of xenograft models. The OT-1 immunocompetent SEAKER models developed in this study provided systems in which to test important pharmacokinetic parameters of SEAKER cells in advance of clinical translation. In the context of a complete immune system, SEAKER cells are still able to localize at high concentrations to antigen-positive tumors and to deliver cargo with minimal off-tumor localization.

Solid tumors present additional challenges to effective adoptive T cell therapy in comparison to hematopoietic cancers. T cell penetration in solid tumors is often aberrant, due to altered tumor extracellular matrix, trapping by suppressive immune cells, and necrotic acellular cores (29,30). Solid tumors also express antigens that are found on vital healthy tissues. This leads to toxicities that limit the effectiveness of many solid tumor therapies. Additionally, solid tumors are notoriously heterogenous. Many tumor-specific antigens are only present in a subset of cells within the tumor, leading to inevitable escape after adoptive T cell therapy. The B16-SIIN tumor model is highly aggressive, with high resistance to T cell-mediated killing *in vivo*. Addition of the SEAKER enzyme/prodrug system led to delayed solid tumor growth and improved survival without the characteristic systemic toxicity of chemotherapeutics, even in the absence of substantial T cell killing.

Immunogenicity is a significant barrier for synthetic or living therapeutics. However, adenovirus and microbial-derived therapies have seen clinical success despite their potential immunogenicity (31,32). Synthetic proteins developed in a given species may still elicit an immune response in the same species, due to small changes in the amino acid sequences, new sequences at fusion junctions of chimeric proteins, or antigen presentation of the synthetic protein. For example, both mice and humans mount humoral immune responses against species-matched CAR proteins (8). These immune responses may explain why some patients do not achieve complete responses when treated with CAR-T cells or have poor persistence or responses to later infusions of the same product. Despite their immunogenicity, CAR-T cells and other synthetic therapeutics are efficacious for some B-cell cancers. Persistence of adoptive T cells, even if immunogenic, may be prolonged by use of effective lymphodepleting regimens. In syngeneic mice, lymphodepleting preconditioning is a requirement for adoptive T cell engraftment. In the current study, both the SEAKER enzymes and cells elicited an immune response in the immunocompetent model, even within the short timeframe of the aggressive B16F10-SIINFEKL model (Figure 6, Supplemental Figure 13). Nonetheless, the SEAKER enzymes remained functional despite antibody binding, and secondary engraftment of SEAKER cells could be achieved through stringent preconditioning regimens.

The SEAKER platform is a modular small-molecule delivery platform that takes advantage of intrinsic T cell function to deliver therapeutic cargo to tumors. Antigen stimulation increases and localizes therapeutic payload to tumor via two mechanisms. Firstly, stimulated T cells are retained in their target tissues and undergo proliferation. This leads to high tumor concentrations of T cells, while trace numbers of cells localize to off-target tumors (Figure 2, Figure 3). Secondly, activated T cells express more retroviral transgenes (33). T cell activation leads to activation of retroviral promoter LTRs and expression of downstream genes (34). Although we used constitutively expressed enzymes, these two mechanisms localize the enzyme to its target tissue. When tumor, SEAKER cells, and prodrug are all injected *IP*, toxicity in the mice is observed, due to systemic leakage of unmasked AMS from the peritoneal cavity. However, in a more clinically relevant adoptive transfer model using *IV* administered SEAKER cells and a solid subcutaneous tumor, anti-tumor benefit was observed without the characteristic toxicity of AMS. The SEAKER platform has the highest therapeutic window when T cells must traffic into the tumor from circulation. Once in the target tissue, any cell in the vicinity of unmasked drug will be susceptible, offering the possibility to kill antigen negative cells and stromal cells. Future iterations of the SEAKER platform may also include T cell activation inducible promoters, such as NFAT, to further control the delivery of payloads.

The potential for targeted delivery of synthetic cargo to tissues has applications outside of cancer. Enzyme–prodrug therapy could be used in a variety of diseases to deliver drugs directly to the affected tissues. In the current study, we expanded the application of SEAKER cells to include both CAR T cells and TCR T cells, demonstrating the wider translational potential of targeted micropharmacies. Cellular micropharmacy-directed therapy could be used in a variety of disease states, such as inflammatory or autoimmune conditions, to deliver potent immunosuppressives to target tissues, sparing systemic immunosuppression. Critically, a barrier to further application of the SEAKER platform in inflammatory disease states is the need for cells that do not kill their target. This will require further engineering and exploration of other cell types as targeted micropharmacies.

## Methods

### Prodrugs

Ceph-AMS and AMS-Glu were synthesized as previously described (7).

### Recombinant proteins

CPG2 and β-Lac proteins were produced and purified by GenScript as previously described (7). Constructs contain C-terminal hemagglutinin (HA) and His6 epitope tags and were purified by nickel affinity chromatography.

### Generation of Retroviral Vectors and Producer Cell lines

To generate murine SEAKER cells, the gene encoding β-lactamase (β-Lac) and carboxypeptidase G2 (CPG2) were cloned into the SFG gamma retroviral vector (generously provided by the R. Brentjens lab, MSKCC) alongside a P2A self-cleaving peptide and green fluorescent protein (GFP) or alongside the CAR constructs for the anti-murine CD19 scFv or the 4H11 anti-MUC16 scFv with murine CD28 and CD3ζ genes. Trackable β-Lac SEAKER cells were generated by cloning the β-Lac enzyme upstream of a *Gaussia* luciferase (gLuc) gene separated by a P2A site, and upstream of the mCherry gene via a T2A site.

Standard molecular biology techniques and Gibson assembly were used to generate all constructs. Retroviral producer cell lines were generated by transiently transfecting of H29 cells with named constructs utilizing standard CaPO_4_ transfection (Promega, Madison, WI, USA). Supernatant from transfected H29 cells was used to transduce Phoenix-ECO (pECO) stable producer lines for murine cells (ATCC, Manassas, Virginia, USA), or Galv9 producer cells for human transduction (generously provided by the Brentjens lab). Stable cell lines were sorted for top expressers.

### Cell Culture

All retroviral producer cell lines, human and murine tumor cells lines, and primary T cells were maintained in RPMI-1640 medium (MSKCC Media Core Facility). Mouse T cells were maintained in RPMI-1640 supplemented with sodium pyruvate and 2-mercaptoethanol (both from Life Technologies Corporation, Carlsbad, CA, USA). All media was supplemented with 10 % fetal bovine serum, 2 mM L-glutamine, 100 IU/mL penicillin, and 100 μg/mL streptomycin. All cells were routinely checked for mycoplasma and treated with Plasmocin (Invivogen, San Diego, California, USA) for two weeks, if positive.

### β-Lac nitrocefin cleavage assay

Nitrocefin assay was completed as previously described (7). Samples in PBS, water, or phenol red-free RPMI-1640 were serially diluted (2-fold) and mixed 1:1 with 0.2 mM nitrocefin (Abcam, Cambridge, UK). Samples were incubated 1–16 h at room temperature and absorbance at 490 nm was read on a SpectraMax M2 plate reader (Molecular Devices, San Jose, California, USA). Data were analyzed with SoftMax Pro software 6.2.2. Recombinant β-Lac standard curves were run simultaneously to quantify β-Lac enzyme concentrations in samples when feasible.

### CPG2 methotrexate cleavage assay

Sample with CPG2 enzyme was incubated with methotrexate at 450 µM final concentration overnight. Absorbance at 320 nm was recorded on a NanoDrop spectrophotometer (Thermo Fisher Scientific, Waltham, Massachusetts, USA), and decrease in UV signal signified substrate cleavage.

### T cell Isolation and Modification

Mouse T cells were isolated from spleens of naïve mice by mechanical disruption using a 100 µm cell strainer. Splenocytes were collected and red blood cells were lysed using ACK lysis buffer to remove red blood cells (Corning, Manassas, VA, USA). Splenocytes were activated overnight with CD3/CD28 Dynabeads (Life Technologies, Carlsbad, California, USA) and 50 IU/mL human IL-2. Activated T cells were transduced by centrifugation with retroviral supernatant from transduced Phoenix Eco cells on RetroNectin (TakaraBio, Kusatsu, Shiga, Japan) -coated plates for 2 consecutive days. Transduction efficiencies ranged between 25% and 75%. A minimum transduction efficiency of 25% was used for all experiments, unless otherwise noted.

### Flow Cytometry

Cell samples were washed and stained in flow buffer (PBS, 1% FBS, 0.1% sodium azide) for 20–30 min. Antibodies used in the study are as follows: APC or FITC anti-mouse CD45.1 (clone: A20, Tonbo Biosciences), Alexa Fluor 700 anti-mouse CD45 (Clone: 30-F11, BioLegend), PE anti-CD3 mouse monoclonal antibody (clone: UCHT1, BioLegend). Data was collected using a Guava easyCyte HT Flow Cytometer (Luminex, Austin, Texas, USA), a LSRFortessa (BD, Franklin Lakes, NJ, USA), or Cytoflex LX (Beckman Coulter, Pasadena, CA). Data were analyzed using Flowjo v10.4 software (Flowjo, Ashland, OR, USA).

### Enzyme-linked immunosorbent assay (ELISA) analysis

Sandwich ELISAs were performed on 96-well Immulon HBX plates (Thermo Fisher Scientific). A mouse IgG anti-β-Lac antibody (clone: 3E11.G3, Thermo Fisher Scientific) was used to capture recombinant β-Lac and primary murine serum samples were used as primary antibody. A polyclonal anti-mouse IgG HRP antibody was used as detection antibody (Novus Biologicals, Littleton, Colorado, USA). Protein was detected using TMB (3,3′,5,5′-tetramet hylbenzidine) substrate (Thermo Fisher Scientific) and H_2_SO_4_ acid quench, and read on a SpectraMax M2 plate reader. Data were analyzed with SoftMax Pro software.

### Cytotoxicity assays

T-cell and prodrug cytotoxicity assays with secreted enzymes were performed after 24 to 48 hours using CellTiter-Glo (Promega, Madison, Wisconsin, USA) or luminescence using firefly (ffLuc) expressing tumor cells. For CellTiter-Glo based experiments, cells were analyzed in duplicate or triplicate wells of a 96-well plate and equivalent volume of CellTiter-Glo reagent was added to each well. Following a 10-min incubation at room temperature, samples were transferred to White 96-well Optiplates (Perkin Elmer, Waltham, Massachusetts, USA) and luminescence was measured on a SpectraMax M2 plate reader. Data were analyzed with SoftMax Pro software. The cytotoxicity of SEAKER cells was determined by luciferase-based assays. EL-4 OVA cells expressing ffLuc were used as target cells. Effector and tumor target cells were co-cultured in triplicate at the indicated E:T (effector-to-target) ratio using clear bottom, white 96-well assay plates (Corning 3903) with 2 x 10^4^ target cells in a total volume of 200 µL. Target cells alone were plated at the same cell density to determine maximum luciferase activity. Cells were cocultured for 4–18 h, at which time D-luciferin substrate (Gold Biotech, St. Louis, Missouri, USA) was added at a final concentration of 0.5 µg/µL to each well. Emitted light was detected in a Wallac EnVision Multilabel reader (Perkin Elmer). Target lysis was determined as (1-(RLUsample)/(RLUmax))x100.

### *In vivo* experiments

All experiments were performed in compliance with all relevant ethical regulations and in accordance with MSKCC IACUC protocol 96-11-044. All mice were included in the analyses and no attrition was noted. Mice were 6 to 12 weeks old and weighed 18-30 grams when treated. Both male and female mice were used. Female mice were randomized into groups to allow balance in groups for tumor growth before treatment. Male mice were maintained with their initial littermates to avoid fighting. Experiments were not blinded, but results were confirmed by blinded third parties. Experiments were replicated 2 or more times as indicated in the legends.All BLI was performed using a Xenogen IVIS Spectrum and analyzed using Living Image software 4.7.4 (Xenogen Biosciences, Cranbury, New Jersey, USA) or Aura 4.0 (Spectral Instruments Imaging, Tucson, Arizona, USA). Methods for supplemental *in vivo* experiments are described in their corresponding figure legends.

#### EL4-OVA peritoneal lymphoma model

C57/BL6 mice were engrafted *IP* with 2 x 10^6^ EL-4 OVA (ATCC) cells on day −6 or −7. Transduced SEAKER cells were engrafted *IP* on day 0. For β-Lac efficacy experiments, 4 mg/kg of Ceph-AMS was administered beginning on day 1, for 3 doses *BID.* For CPG2 efficacy experiments, 50 mg/kg AMS-glu was injected *IP* from day 1 to day 5 *BID*.

#### B16F10 SIINFEKL melanoma models

C57/BL6 mice were engrafted *SC* with 1 x 10^6^ B16F10 melanoma cells engineered to express the SIINFEKL peptide (B16-SIIN) on day –7 (generously provided by the Andrea Schietinger Lab, MSKCC). Tumors were engrafted using a 1:1 mixture with matrigel. On day –1, mice were treated *IP* with 100 mg/kg cyclophosphamide (Sigma-Aldrich, St. Louis, Missouri). On day 0, mice were engrafted *IV* via retro-orbital injection with 1–3 x 10^6^ bulk murine T cells.

#### B16F10 SIINFEKL OT-1 T cell Kinetics and localization

Experimental parameters described in "B16F10 SIINFEKL melanoma models" above were used. Tumors were harvested at days 2, 5, 7, 10, 14 by mixing cut tumors with Mouse Tumor Dissociation Kit (Miltenyi Biotec, Bergisch Gladbach, North Rhine-Westphalia, Germany) in RPMI-1640 and running on a gentleMACs (Miltenyi Biotec) for 10–20 min. Spleen samples were ground through a 100uM cell strainer into ACK lysing buffer and washed. Single cell suspensions were washed and prepared for flow cytometry as described in "Flow Cytometry" above.

For BLI T cell tracking, mice were imaged on days 1, 2, 5, 7, 10, and 14 by injection of 100 µg of coelenterazine per mouse via retro-orbital *IV* injection, with an exposure of 60 seconds.

For lymphoid organ SEAKER cell experiments, mice were harvested on day 6. Tumor and spleens were harvested as described above. Inguinal lymph nodes and lungs were removed and mechanically dissociated through a 100 µm strainer into RPMI-1640. Bone marrow was collected by removing femurs and crushing them with mortar and pestle in RPMI-1640. Blood samples were collected through cheek bleeds into heparinized tubes. ACK lysing buffer was used to lyse blood cells for 5 minutes. Collected cells were washed in PBS and prepared for flow cytometry as described.

#### B16F10 SIINFEKL OT-1 β-Lac Enzyme Activity Model

Experimental parameters described in "B16F10 SIINFEKL melanoma models’’ above were used. On day 5 post T cell engraftment, tumor samples were harvested through a cell strainer into PBS to solubilize the cell-free proteins. Samples were centrifuged at 12,000g to remove all debris and clarified supernatant fluids were run in the nitrocefin assay.

#### B16F10 SIINFEKL OT-1 Efficacy Model

Experimental parameters described in "B16F10 SIINFEKL melanoma models’’ above were used. Prodrug injections spanned days 4–8, with 4 mg/kg Ceph-AMS *IP, BID*. Tumor length and width was measured and tumor volume was determined using the formula: [(LxW)^2]/2. Mice were euthanized once tumor volume exceeded 2 cm^3^ or if excessive distress or tumor ulceration was observed.

### Statistical Analysis

Data reported as mean ± SD unless otherwise noted. For *in vitro* experiments, technical replicates are displayed. For *in vivo* and *ex vivo* experiments, individual mice are displayed. Log-rank, unpaired or paired *t* tests were performed using Prism 8 software (GraphPad, La Jolla, CA, USA) when appropriate. Statistical significance was indicated as: * = *p* < 0.05, ** = *p* < 0.01, *** = *p* < 0.001, **** = *p* < 0.0001.

### Contributions of authors statement

C.M.B., P.W., M.D., K.V., B.C.C., M.A.B., J.R.P., A.C., and K.G.K., performed experiments and analyzed the data; C.M.B., P.W., M.D., K.V., B.C.C., M.A.B., J.R.P., A.C., K.G.K., D.S.T., D.A.S., designed the experiments; C.M.B. and P.W., wrote the initial manuscript draft; D.S.T. and D.A.S. supervised the project, provided funding, and edited the manuscript; and all authors reviewed and contributed to the manuscript.

## Acknowledgements

We thank the Renier Brentjens lab (MSK) for providing reagents and expertise throughout this project.

Financial support for this work was provided by the Leukemia and Lymphoma Society (to D.A.S.), grants from the National Institutes of Health (P30 CA008748 to C. B. Thompson, R01 CA055349 to D.A.S., P01 CA023766 to D.A.S. and D.S.T., R35 CA241894 to D.A.S. and D.S.T., F31 CA254311 to C.M.B., T32 CA062948–Gudas to B.C.C., and F31 CA261179 to B.C.C.), Tudor Funds (to D.A.S.), Commonwealth Foundation and Experimental Therapeutics Center of MSKCC (to D.S.T. and D.A.S.), and the Memorial Sloan Kettering Maximizing Excellence in Research, Innovation and Technology Sawyers Fellowship (to C.M.B.).

## Supplemental for: Host-cell Interactions of Engineered T cell Micropharmacies

**Supplemental Figure 1.**
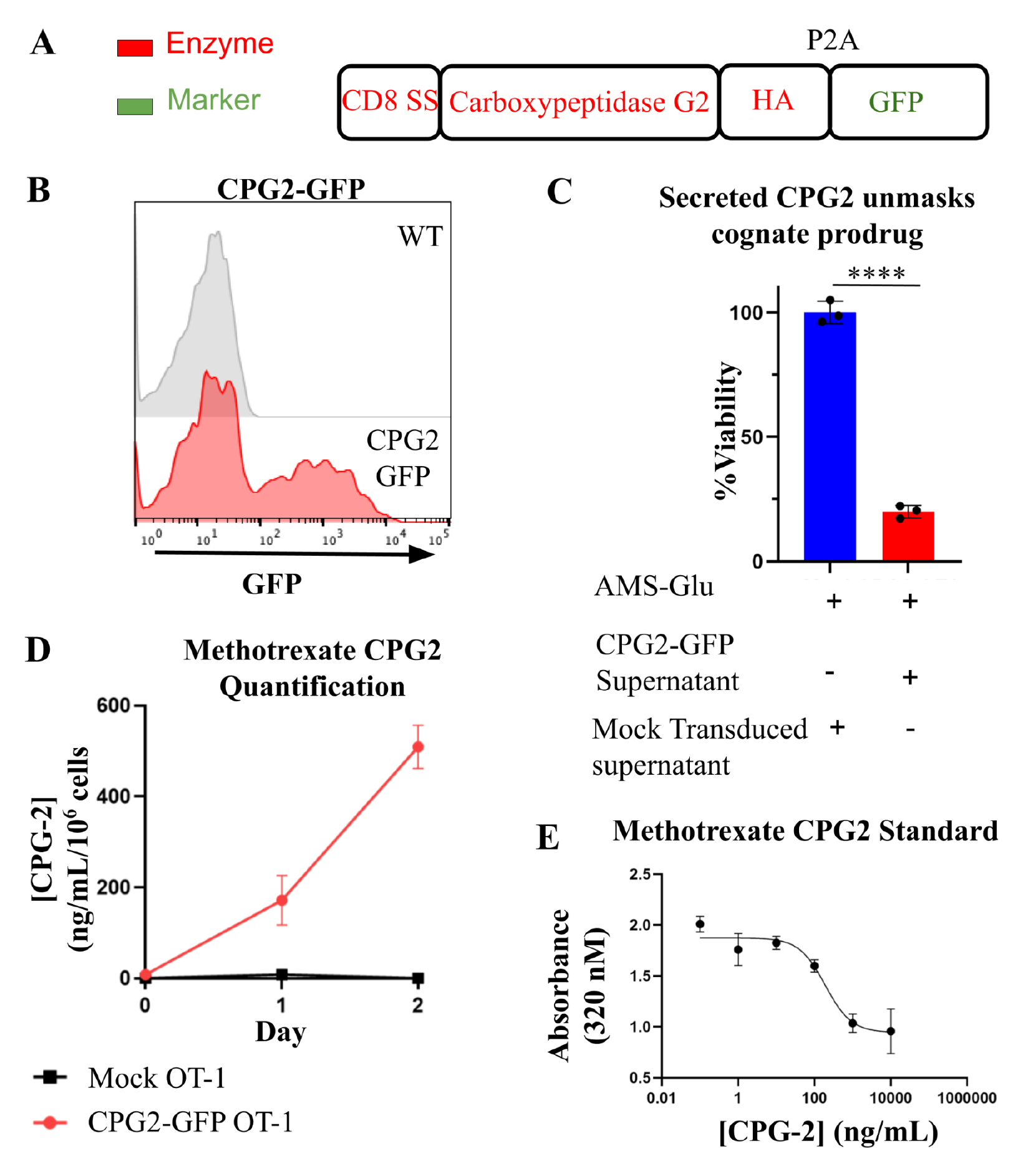
OT-1 syngeneic CPG2 SEAKER cells secrete functional enzyme and synergize with prodrug to kill cancer cells. (A) Schematic of SEAKER enzyme secretion cassette for carboxypeptidase G2 (CPG2) with a CD8 signal sequence and a hemagglutinin (HA) tag. (B) Representative flow histogram of murine primary T cell transduction with the CPG2-GFP or β-Lac-GFP plasmid. (C) Supernatant fluid was collected from primary murine T cells transduced with CPG2-GFP or GFP-Luciferase and B16F10 cells were incubated with 100 uM of the cognate prodrug (AMS-GLU), with or without the indicated supernatant fluid. Cell viability was assessed by CellTiter-Glo. (D) Supernatants from mock or CPG2 secreting OT-1 T cells were quantitated for CPG2 using methotrexate and a standard curve. (E) Representative standard curve of recombinant CPG2 with methotrexate.. * = *p*<0.05; ** = *p*<0.01; *** *p*< 0.001; **** = p<0.0001

**Supplemental Figure 2.**
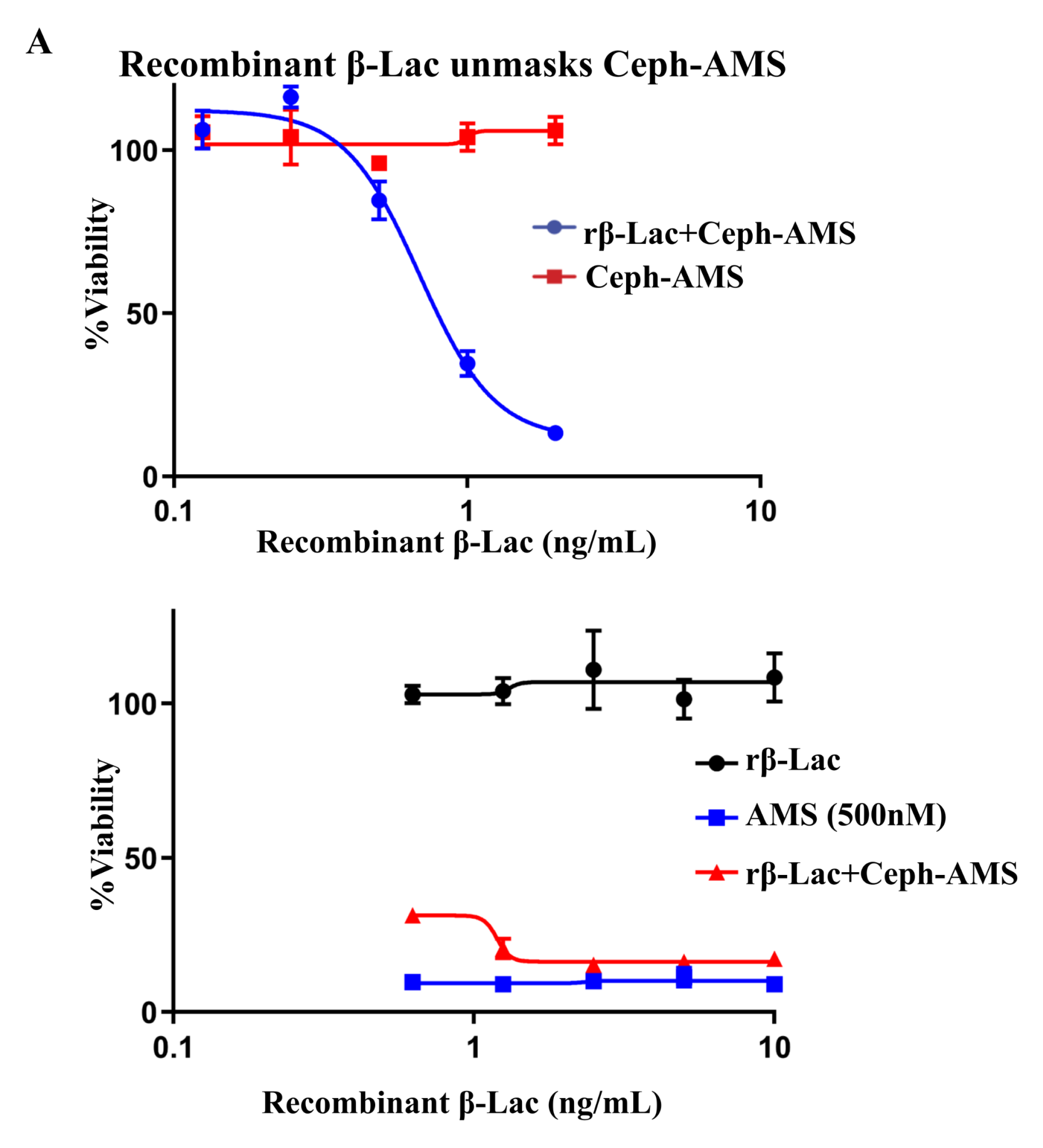
Recombinant β-Lactamase unmasks Ceph-AMS to release AMS drug. Set-2 cancer cells were incubated with 500 nM Ceph-AMS and indicated concentrations of recombinant β-Lactamase, and viability of Set-2 cells were assessed by Cell-Titer Glo after 48 hours.

**Supplemental Figure 3.**
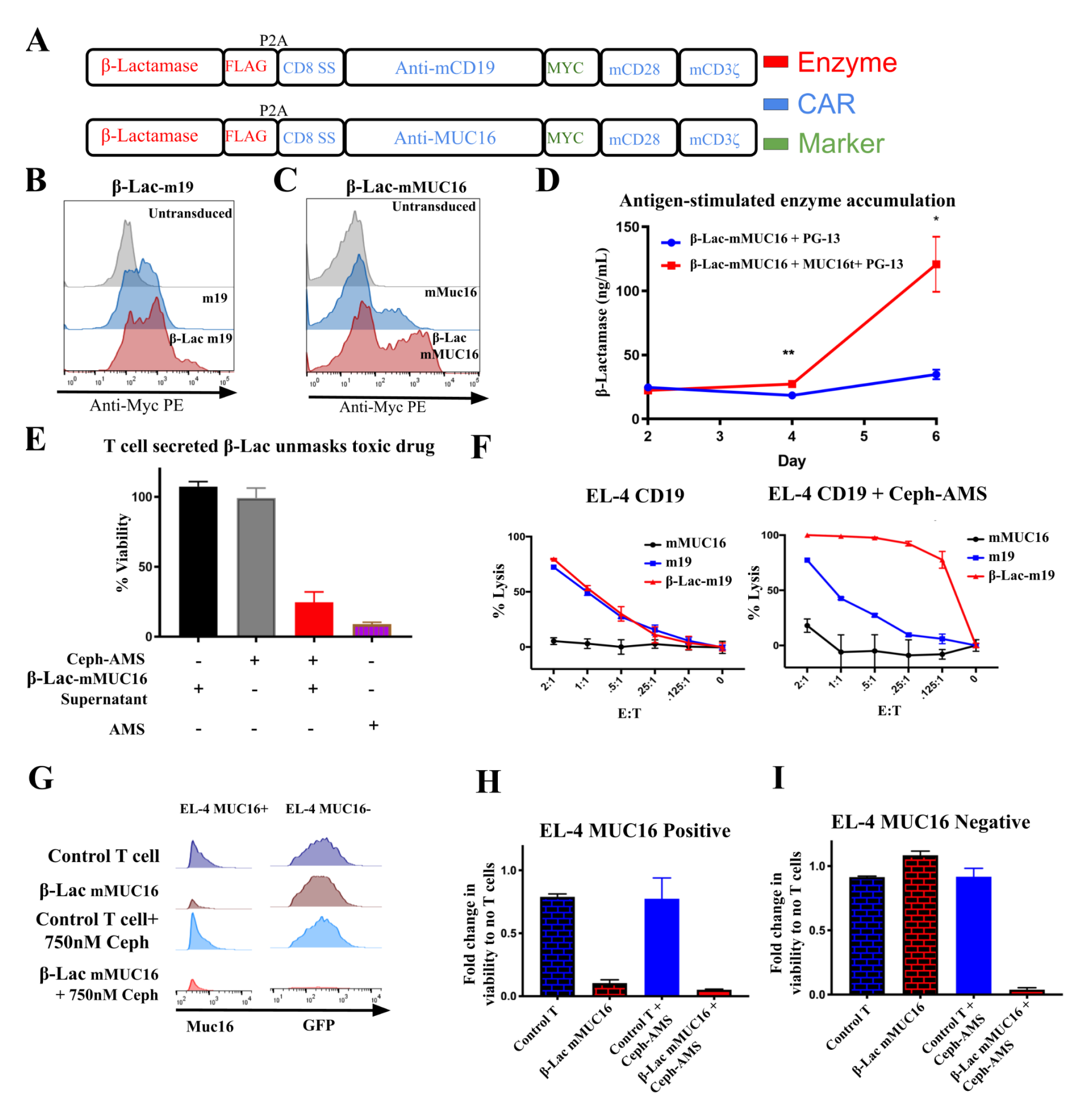
Secreted β-Lac can be integrated with various antigen targeting moieties expressed in murine T cells. (A) Secreted β-Lac SEAKER constructs with anti-murine CD19 (top) or anti-MUC16 (bottom) CARs and myc tags. The CAR signaling domains include murine CD28 and murine CD3ζ. (B) β-Lac-m19 and (C) β-Lac-mMUC16 transduction of primary murine T cells. (D) 1 x 10^6^ β-Lac-mMUC16 SEAKERs were co-cultured with MUC16 positive or negative PG-13 cells. Enzyme secretion was quantified using the β-Lac substrate nitrocefin compared to a standard curve using recombinant β-Lac. (E) ID8 ovarian cancer cells were incubated as indicated with 500 nM Ceph-AMS prodrug, β-Lac-mMUC16 cell supernatant, or 500 nM AMS parent drug for 48 h and viability was determined by luminescence. (F) β-Lac-m19 or wild-type m19 cells were incubated at various concentrations with 2 x 10^4^ EL-4 CD19+ cells for 24 h, with or without 500 nM Ceph-AMS. Viability was assessed by luminescence compared to untreated cells. (G) β-Lac-mMUC16 cells were co-cultured with a 10% MUC16+ and 90% MUC16-EL-4 cell mixture, with or without 750 nM Ceph-AMS for 24 h. MUC16– and MUC16+ cell populations were assessed by flow cytometry. (H) Quantification of panel G, where total count per MUC16– cell populations were divided by untreated samples to determine percent viability. (I) Quantification of panel G for MUC16+ cell populations. Some of the anti-mMUC16 SEAKER data presented here was also published as a supplemental figure in Gardner et. al. 2022(7).

**Supplemental Figure 4.**
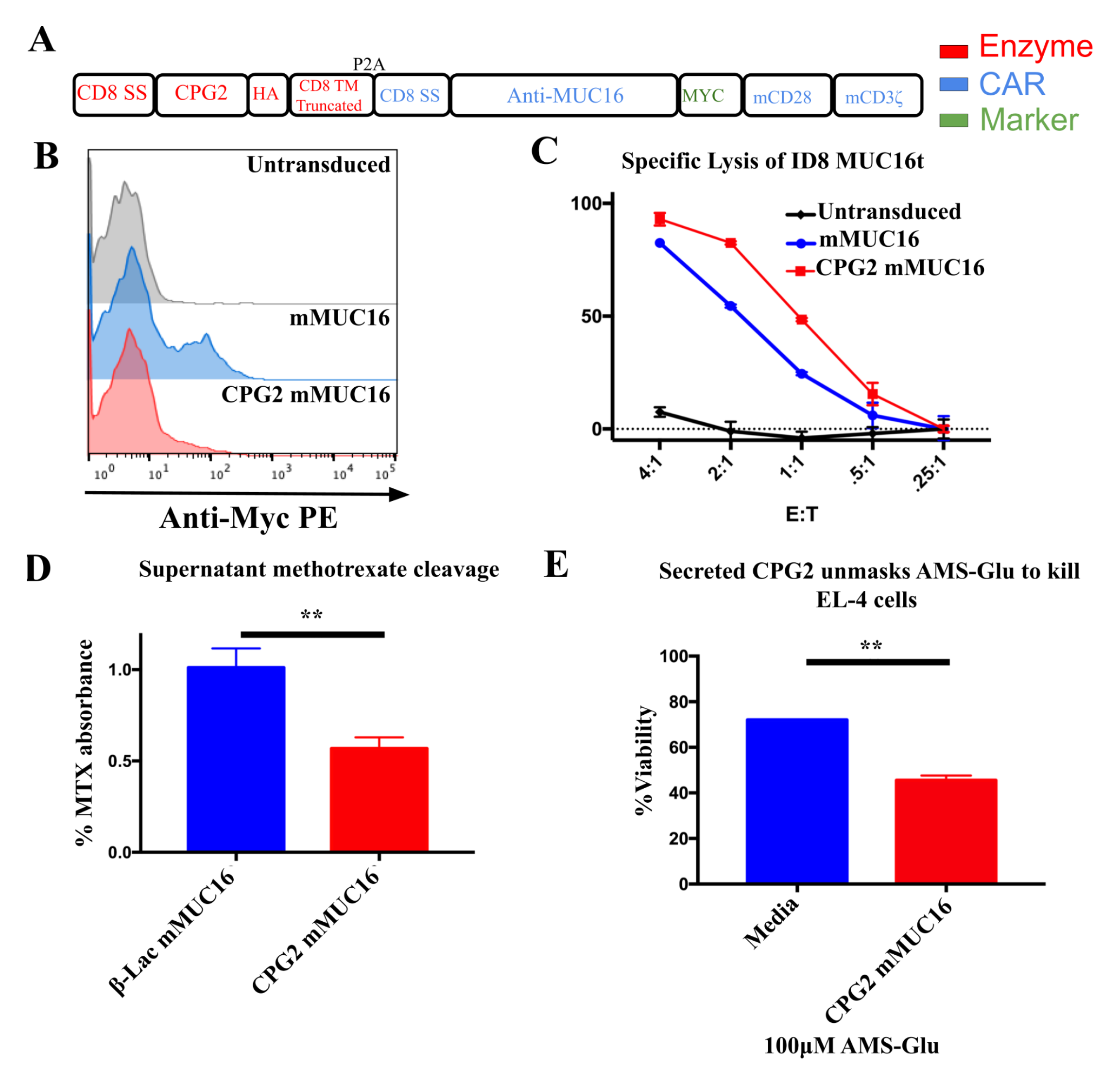
Carboxypeptidase G2 secretion by anti-MUC16 CAR-T cells synergizes with Glu-AMS prodrug to kill cancer cells. (A) Secreted CPG2 SEAKER construct with anti-MUC16 CAR. CAR signaling domains include murine CD28 and murine CD3ζ. (B) CPG2-mMUC16 transduction efficiency of primary murine T cells. This CPG2 construct has a lower transduction efficiency and requires more optimization. (C) Cytotoxicity of wild-type anti-mMUC16 and CPG2-mMUC16 SEAKERs compared using a ID8 MUC16+ cell line as target. (D) Supernatant fluid from CPG2 mMUC16 cells was mixed with the CPG2 substrate methotrexate, and loss of methotrexate absorbance was measured. β-Lac-mMUC16 supernatant fluid was used as a negative control that should not cleave methotrexate. (E) Supernatant fluid from CPG2-mMUC16 cells was mixed with 100 µM AMS-Glu and cultured with EL-4 cells for 48 h.

**Supplemental Figure 5.**
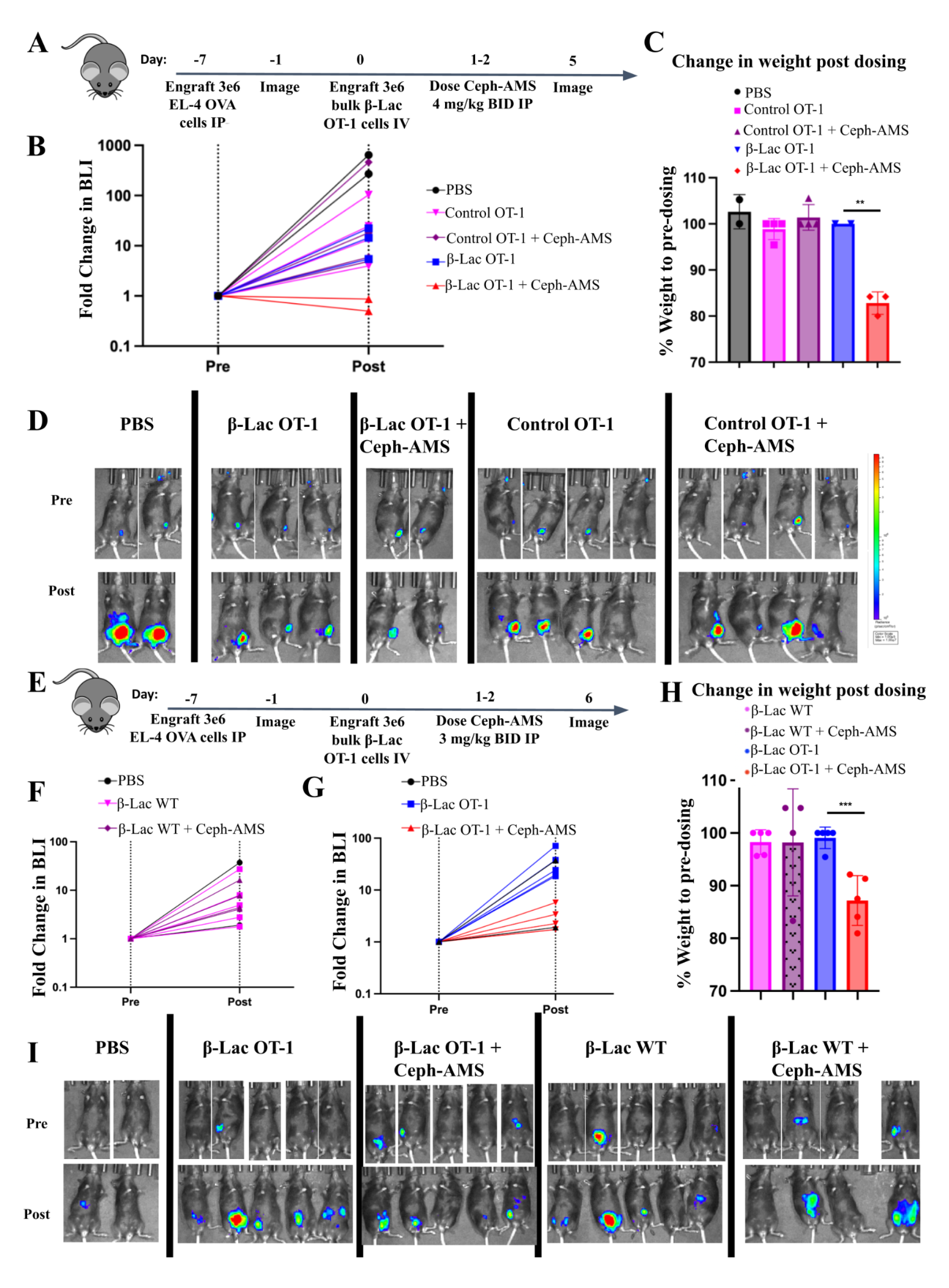
Extended data for figure 1E-F. β-Lac OT-1 SEAKER cells unmask Ceph-AMS prodrug in a peritoneal tumor model. (A) Schematics of intraperitoneal proof-of-concept efficacy syngeneic model. EL-4 OVA cells were engrafted *IP* on day −7. BLI imaging was performed one day before T cell engraftment *IP*. Ceph-AMS was given at 4 mg/kg *IP* for three consecutive doses on days 1 and 2 post T cell engraftment. Imaging was performed on day 5. (B) Graph of fold change in tumor BLI pre and post drugging for experiment (A). Two mice in the combination group died from toxicity. (C) Quantification of change in weight pre and post dosing for experiment (A). (D) BLI imaging of pre and post dosing for experiment (A). (E) Replicate experiment at a lower Ceph-AMS dose. Mice were treated as in (A) using 3 mg/kg Ceph-AMS, and imaged on day 6. (F) Quantification of fold change in BLI of the control WT T cells secreting β-lac for experiment (E). (G) Quantification of fold change in BLI of β-lac OT-1 T cells for experiment in (E). (H) Quantification of change in weight pre and post dosing for experiment (E). (I) BLI imaging of pre and post dosing for experiment (E).

**Supplemental Figure 6.**
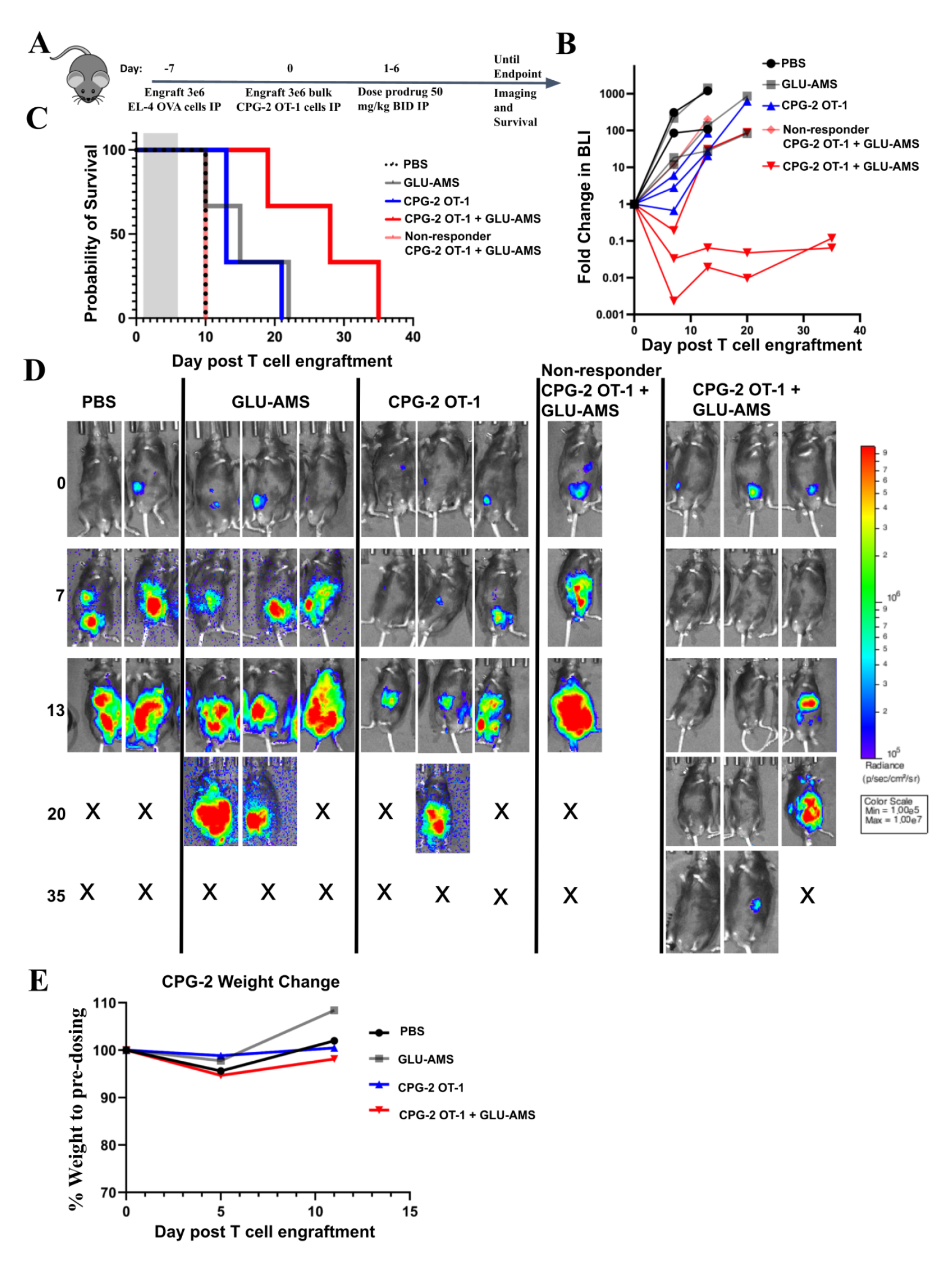
CPG2 OT-1 cells unmask AMS-glu prodrug to delay tumor growth *in vivo*. (A) Schematic for testing CPG2 OT-1 SEAKER cells. 3 x 10^6^ EL-4 OVA cells were engrafted *IP* at day −7. Mice were imaged and engrafted with bulk 3 x 10^6^ OT-1 cells secreting CPG2 on day 0. Mice were given 50 mg/kg *IP*, *BID* from days 1-6. Images were taken of day 7. One mouse that received the CPG combination treatment did not respond to the initial T cell engraftment, and was moved to a non-responder group. This mouse died at the same time as the mice in the PBS treated group. (B) Fold change in BLI for experiment (A). (C) Survival of experiment (A). (D) BLI imaging of experiment (A) An X indicates mouse death. (E) Mean weight values in grams for treatment groups in experiment (A).

**Supplemental Figure 7.**
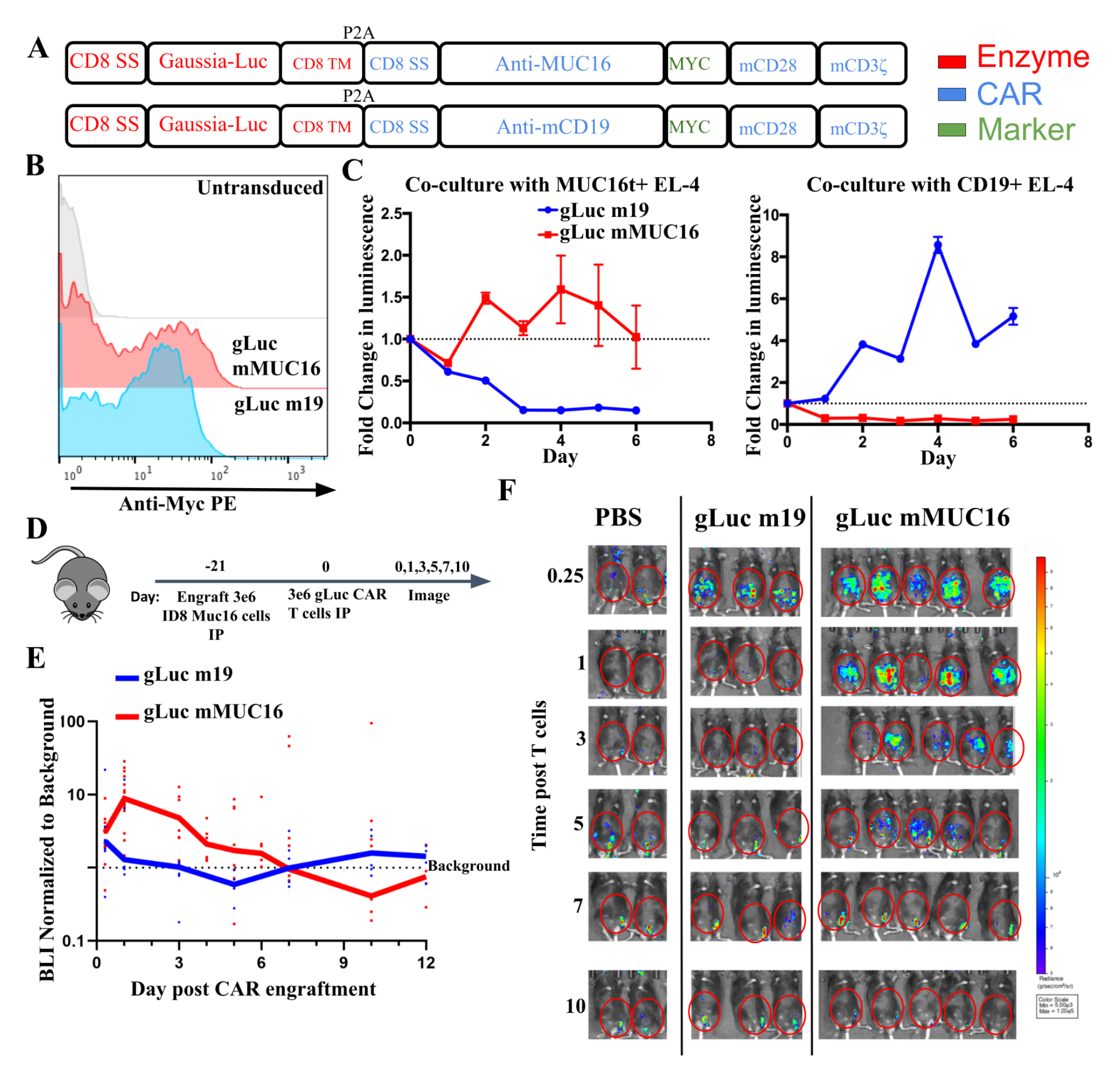
Gaussia luciferase enables tracking of murine T cells in a syngeneic mouse model. (A) Schematic of gaussia luciferase (gLuc) constructs alongside the anti-MUC16 CAR (top) or anti-murine CD19 (bottom) with murine CD28 and murine CD3ζ. (B) Transduction of primary murine T cells with gLuc labeled CARs. (C) gLuc CAR were cocultured with EL-4 cells expressing cognate antigen as indicated, in triplicate. (D) Schematic for gLuc tracking in a peritoneal ovarian tumor model. C57BL/6 mice received 3 x 10^6^ ID8 cells *IP* at day −21. On day 0, 3 x 10^6^ of indicated gLuc CAR T cells were injected *IP* with 100 µg of coelenterazine per mouse, with an exposure of 60 seconds. Images were taken serially at indicated days. (E) Graph of background normalized BLI for the on-target gLuc-mMUC16 (N=17) and off-target gLuc-m19 (N=7) CARs. (F) Representative images of experiment in D.

**Supplemental Figure 8.**
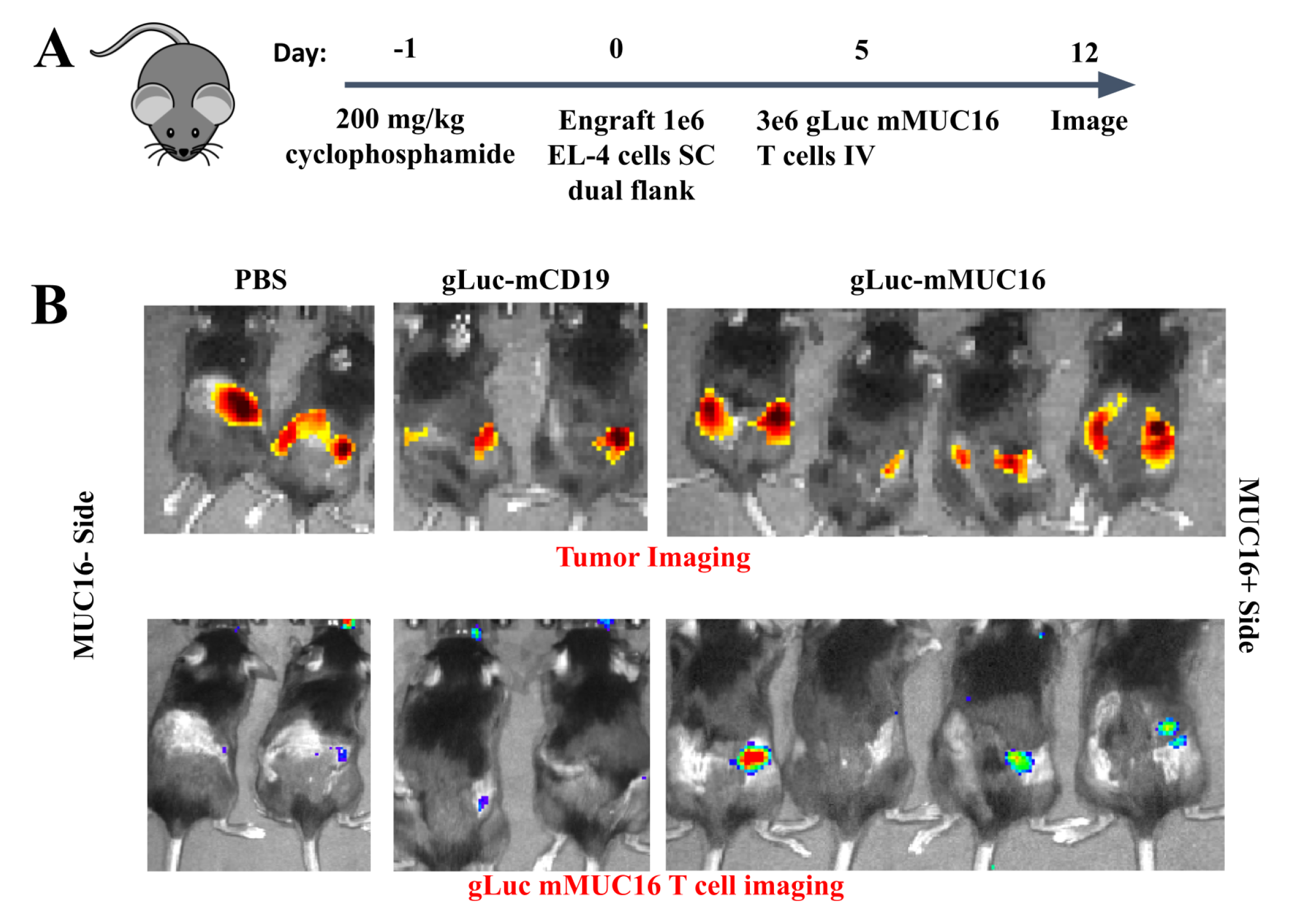
Anti-MUC16 CAR T cells home to antigen positive tumors. (A) Schematic where C57BL/6 mice were preconditioned with cyclophosphamide, followed by a subcutaneous engraftment of 1 x 10^6^ EL4 MUC16- (left flank of mice) or MUC16+ (right flank of mice) tumors. At day 5, 3 x 10^6^ gLuc-mMUC16 cells were engrafted and imaged on day 12. (B) Images of experiment (A). Mice are imaged prone, so the right flank is depicted on the right hand side.

**Supplemental Figure 9.**
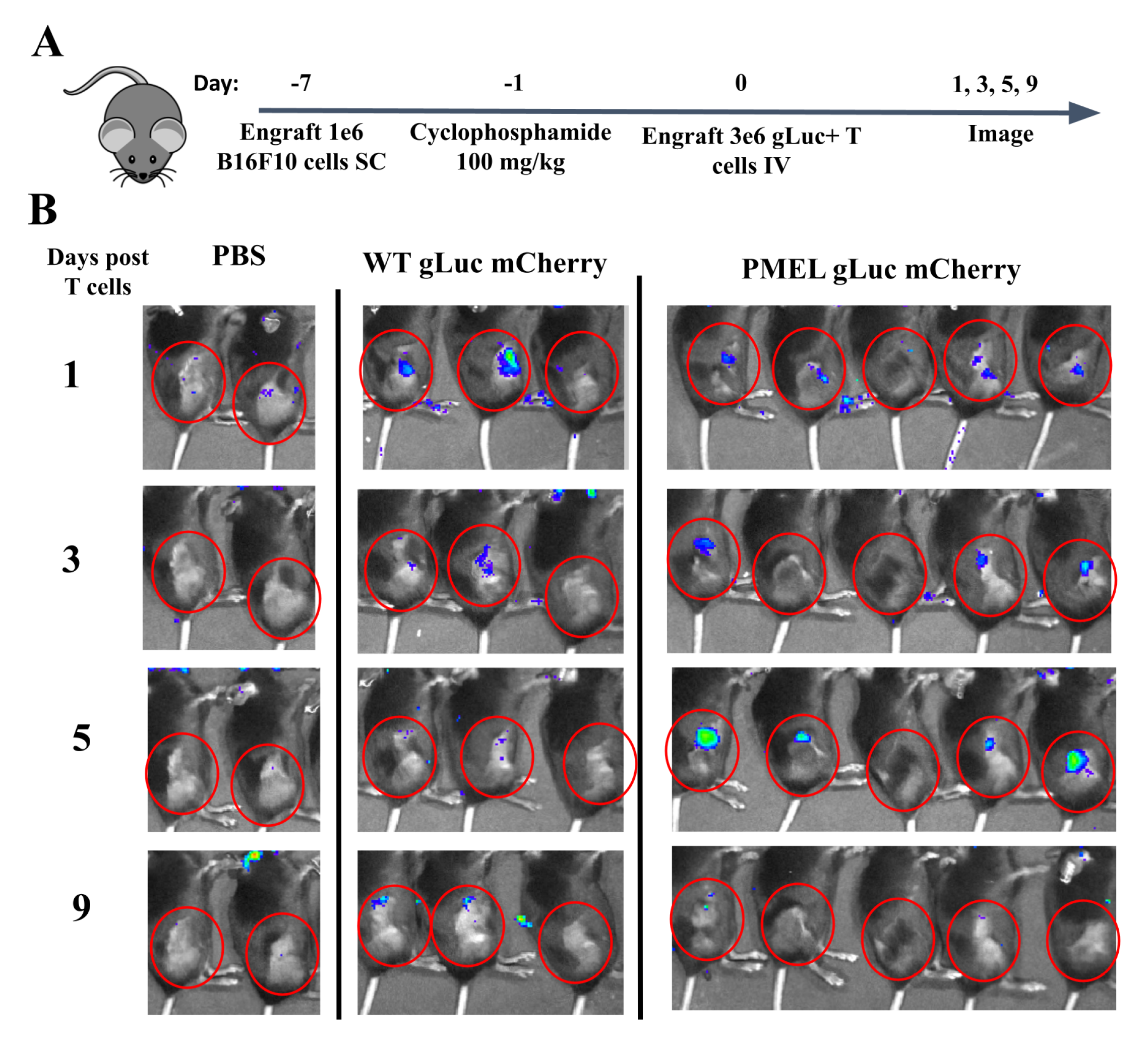
PMEL T cells home to gp100 antigen positive B16F10 tumors. (A) Schematic where C57BL/6 mice received *SC* engraftment of 1 x 10^6^ B16F10 cells at day −7. Cyclophosphamide was injected *IP* at day −1. PMEL gLuc T cells were then engrafted and T cells were imaged serially through injection of 100ug coelentrazine. (B) Images of experiment (A). Due to retro-orbital injection, non-specific signal may be noted near the eyes.

**Supplemental Figure 10.**
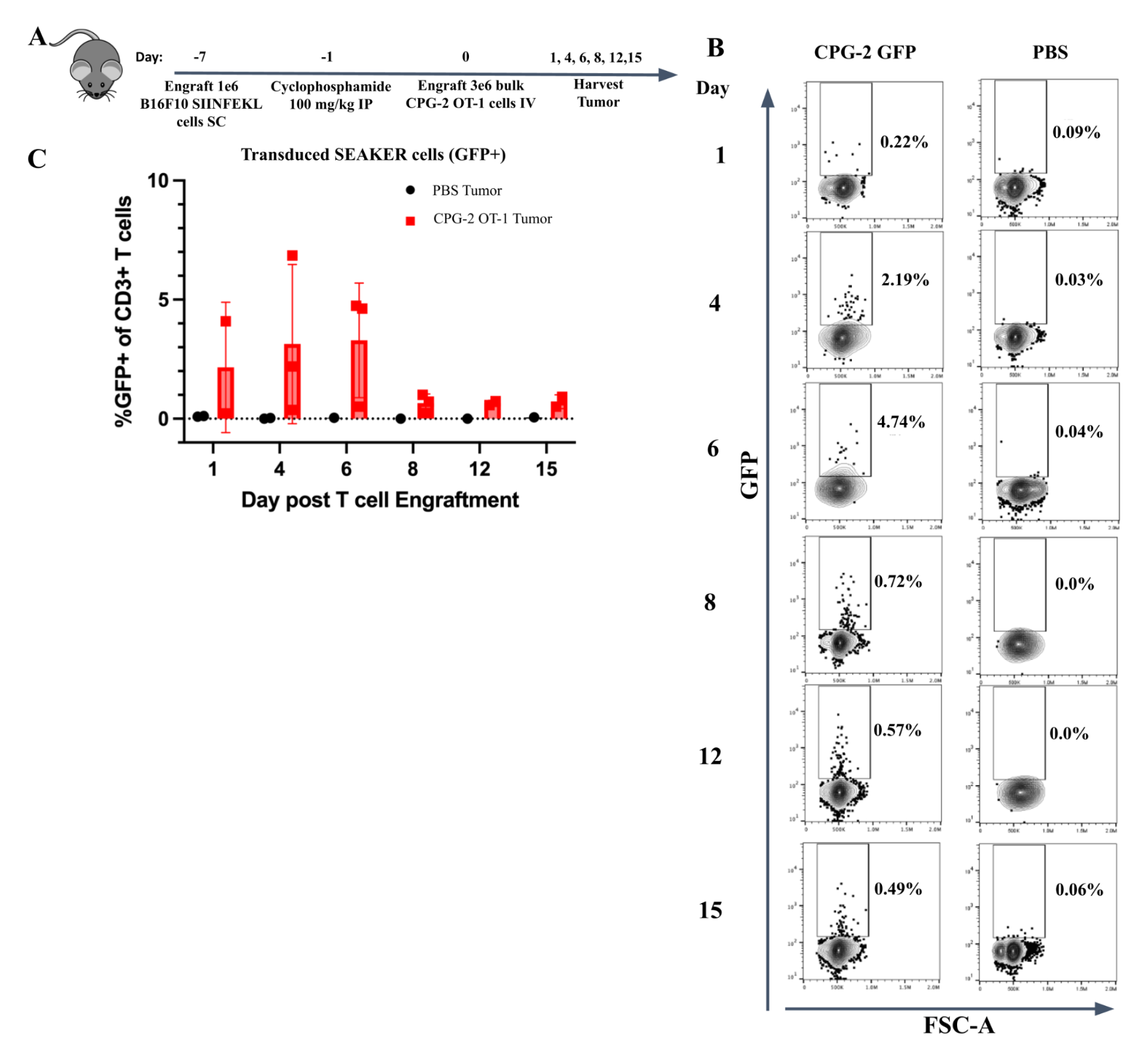
CPG2 OT-1 T cells home to antigen positive B16F10 SIINFEKL tumors. (A) Schematic where C57BL/6 mice received *SC* engraftment of 1 x 10^6^ B16F10 SIINFEKL cells at day −7. 100 mg/kg cyclophosphamide was injected *IP* at day −1. CPG2 OT-1 T cells were then engrafted and tumors were harvested serially at indicated time point. (B) Representative flow images of GFP+ cells of all CD3+ cells. (C) Graph of (B) across all replicates.

**Supplemental Figure 11.**
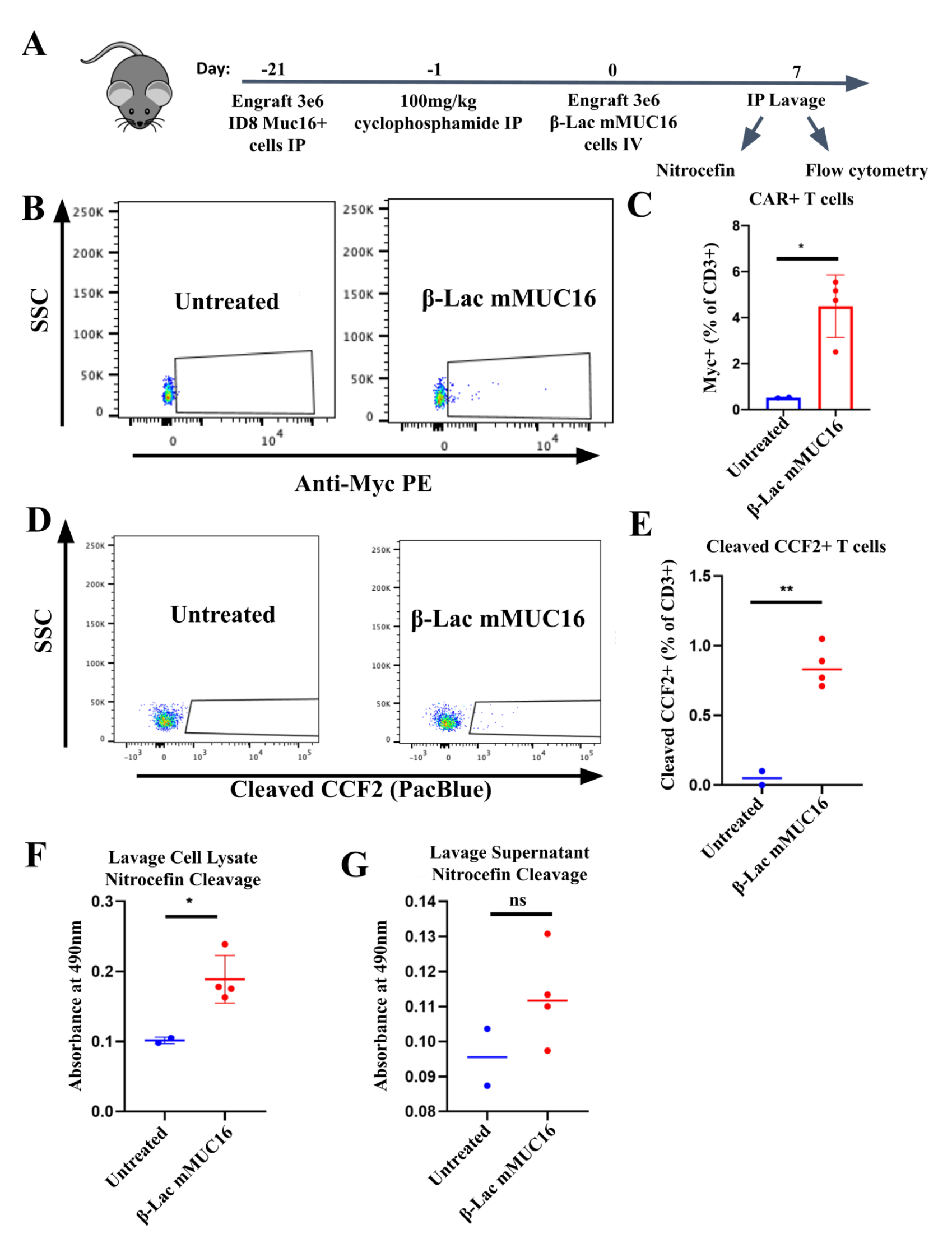
β-lac-mMUC16 SEAKER cells deliver β-Lac to peritoneal ovarian tumors. (A) C57BL/6 mice were engrafted with 3 x 10^6^ ID8 ovarian tumor cells *IP* at day −21. Mice were preconditioned with 100 mg/kg cyclophosphamide at day −1. On day 0, 3 x 10^6^ β-Lac-mMUC16 SEAKERs were engrafted retro-orbitally. Peritoneal lavages were performed on day 7 for flow cytometry and nitrocefin enzyme activity. (B) Flow cytometry of myc staining for CAR+ T cells. (C) Quantification of (B). (D) Flow cytometry of cleaved CCF2 staining, indicating β-Lac activity. (E) Quantification of (D). (F) Cell pellets from peritoneal lavages were lysed and mixed with nitrocefin. A graph of raw absorbance at 490 nm is displayed. (G) Supernatant from the peritoneal lavages were mixed with nitrocefin and raw absorbance at 490 nm was measured.

**Supplemental Figure 12.**
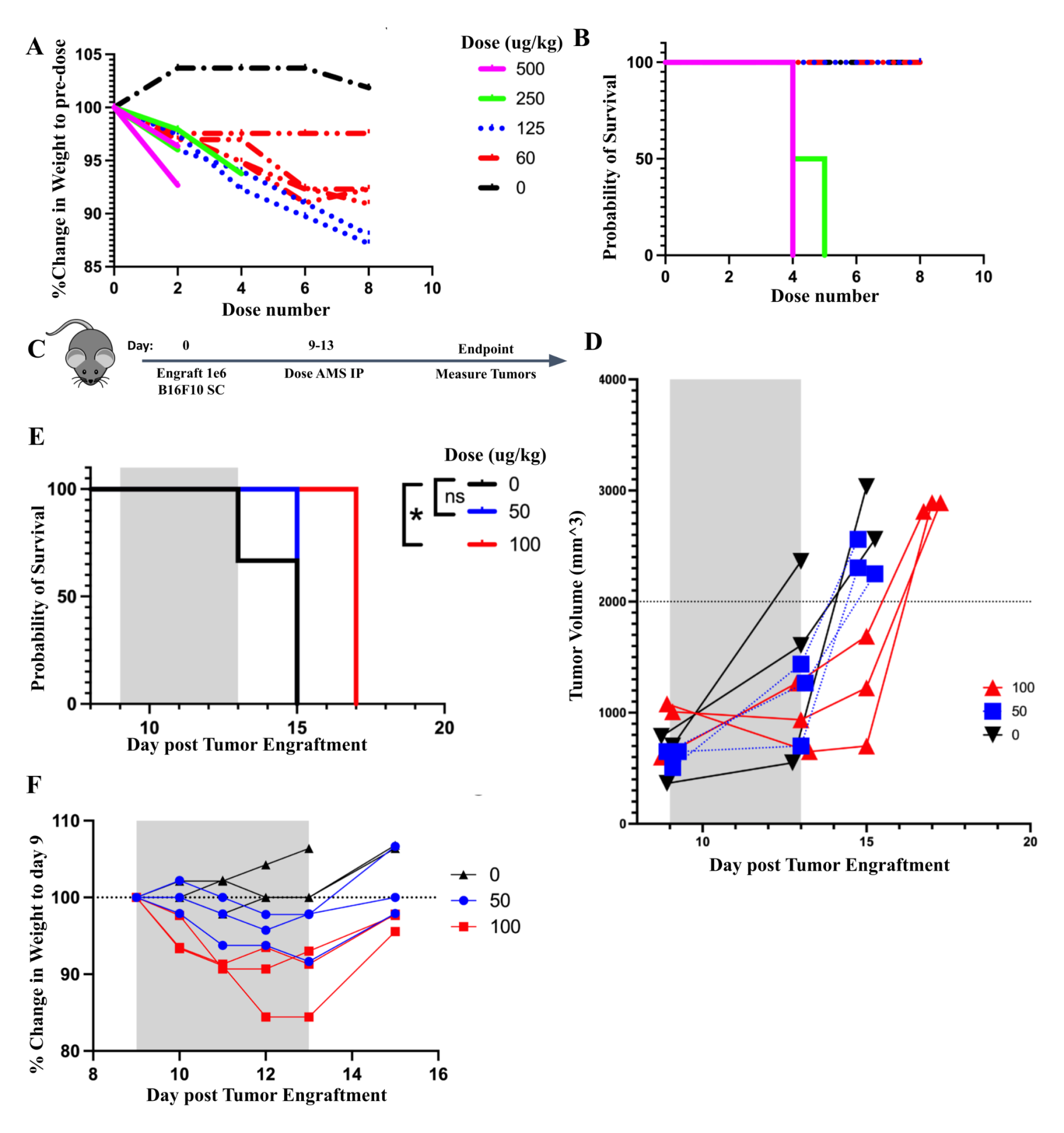
AMS is toxic at therapeutic doses. (A) A dose titration of AMS was performed on C57BL/6 mice. A graph of % change in weight is displayed for each dose (µg/kg). (B) Survival of mice treated with indicated concentrations of AMS (µg/kg). (C) Schematic showing mice engrafted with 1 x 10^6^ B16F10 cells *SC* followed by *BID* dosing of AMS. (D) Tumor measurements of experiment in (C). (E) Survival of mice from experiment in (C). (F) Percent change in weight for mice from experiment in (C).

**Supplemental Figure 13.**
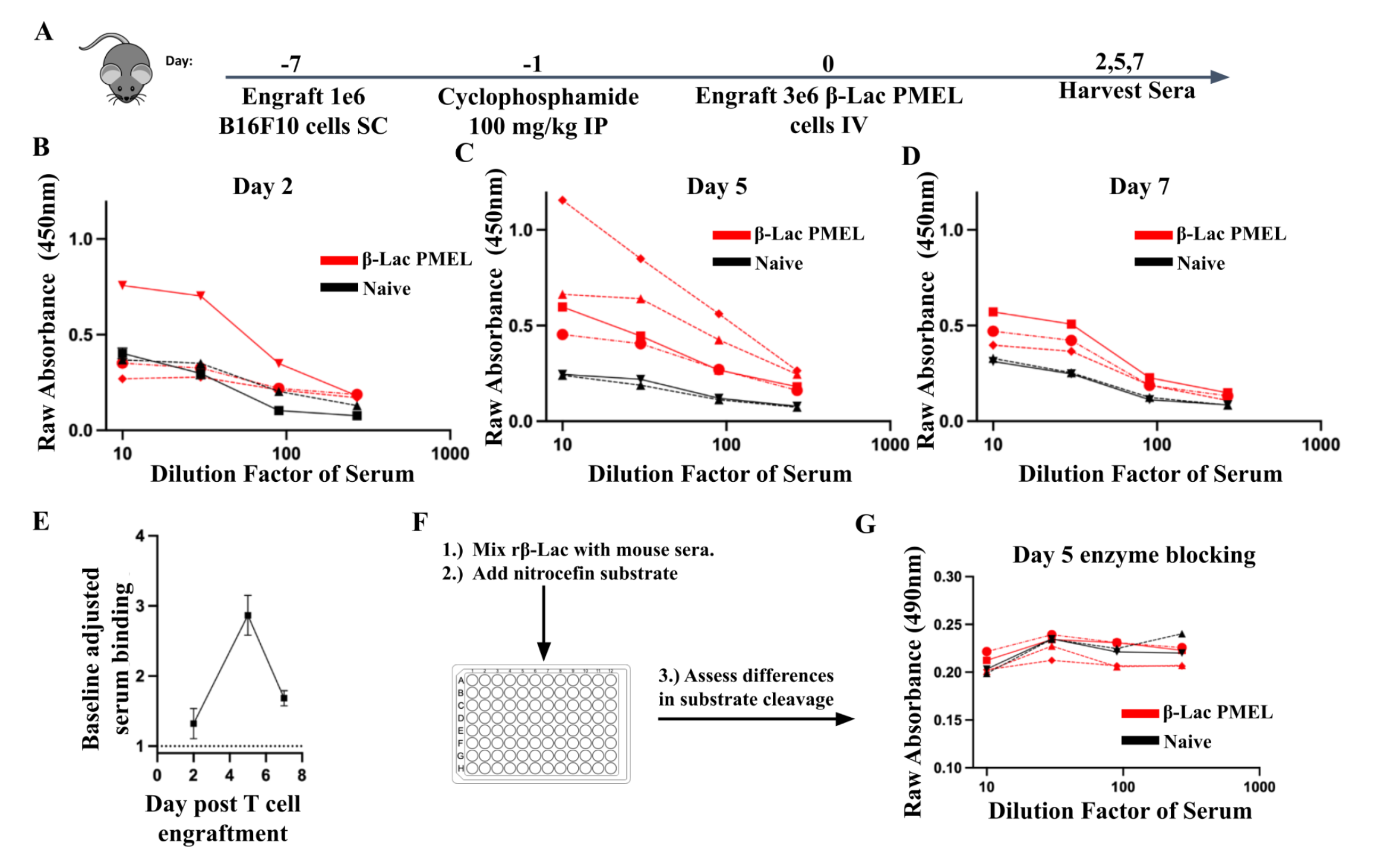
β-Lac SEAKER cells elicit a humoral immune response in C57BL/6 mice. (A) Mice were engrafted with 1 x 10^6^ B16F10 cells *SC* at day −7, 100 mg/kg of cyclophosphamide on day −1, and β-Lac PMEL T cells on day 0. Sera were collected at indicated time points. (B-D) Serum samples were tested for reactivity to recombinant untagged β-Lac at day 2 (B), day 5 (C), and day 7 (D). Raw absorbance is depicted at 450 nm. (E) Average serum binding was normalized to naive mice at the 1:30 serum dilution. (F) Schematic depicting β-Lac enzyme functional assay in which recombinant β-Lac was mixed with serum samples and nitrocefin was added to assess change in activity. (G) Graph depicting raw absorbance values for recombinant β-Lac incubated with indicated serum samples and concentrations.

**Supplemental Figure 14.**
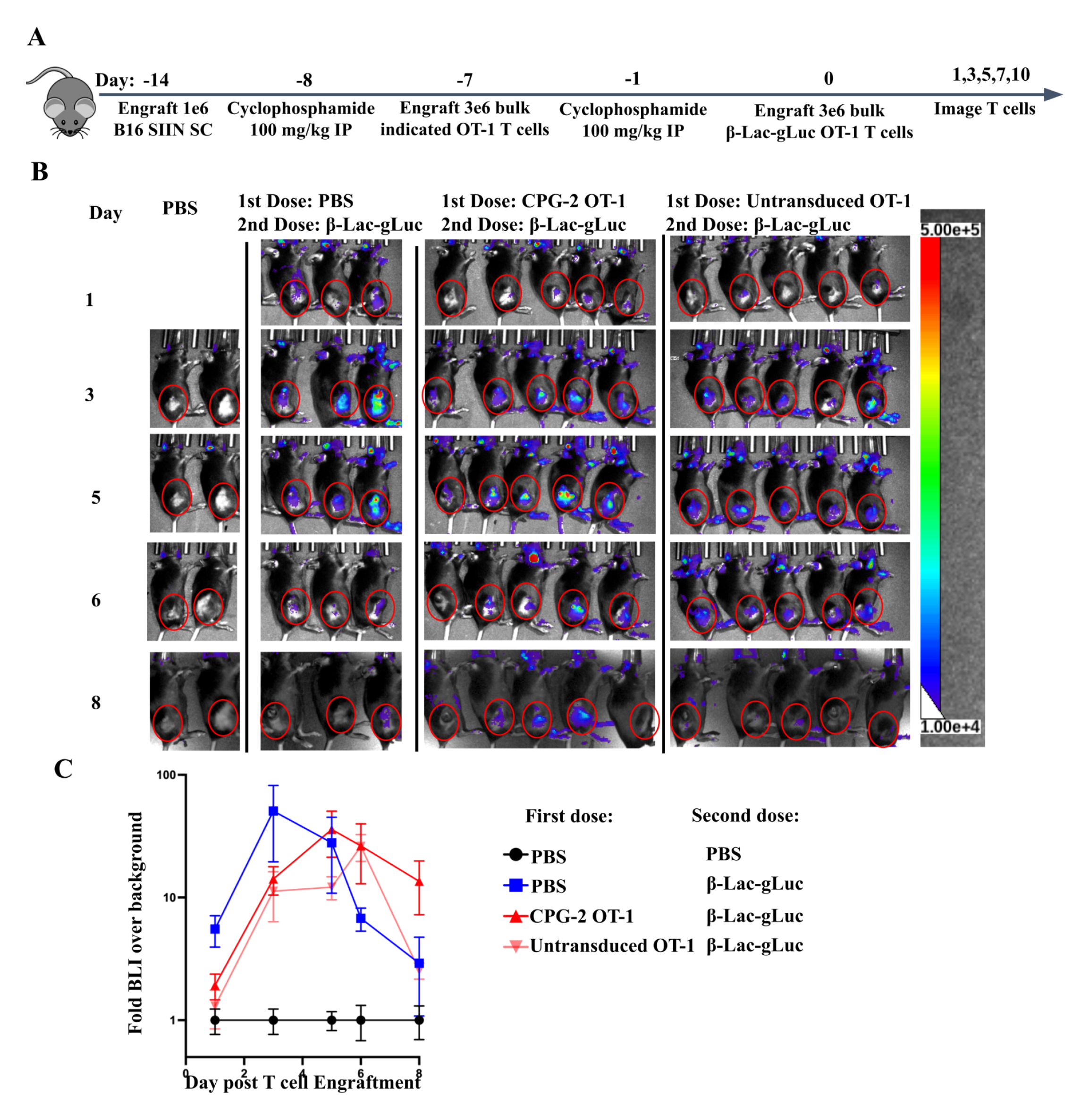
β-Lac SEAKER cells can be re-engrafted with CPG2 SEAKER cells. (A) C57BL/6 mice were engrafted with 1 x 10^6^ B16F10 SIINFEKL cells *SC* at day −14, pretreated with 100 mg/kg cyclophosphamide at day −8 and treated with 3 x 10^6^ indicated OT-1 cells retro-orbitally on day −7. Mice were retreated with 100 mg/kg cyclophosphamide on day −1 and re-engrafted with trackable gLuc+ β-Lac SEAKER cells on day 0. Mice were imaged serially. (B) T cell bioluminescent imaging from experiment (A). (C) Quantification of background adjusted BLI values from (B).

